# Integration of ATAC and RNA-sequencing identifies chromatin and transcriptomic signatures in classical and non-classical zebrafish osteoblasts and indicates mechanisms of *entpd5a* regulation

**DOI:** 10.1101/2024.06.24.600453

**Authors:** Kleio Petratou, Martin Stehling, Ferenc Müller, Stefan Schulte-Merker

## Abstract

Two types of osteoblasts are required to assemble the zebrafish embryonic skeleton: classical osteoblasts homologous to the mammalian cell, and notochord sheath cells, which serve as non-classical osteoblasts. The gene *entpd5a* is critically required for ossification via both types of osteoblasts. Despite the indispensability of zebrafish models in vertebrate research, the genetic regulation of bone formation, as well as mechanisms of transcriptional control of *entpd5a*, remain largely unknown. Here, using a newly generated transgenic line, we isolate classical and non-classical osteoblasts from zebrafish embryos and performed both ATAC-seq and RNA-seq. We analysed results independently and integratively to understand those chromatin dynamics and accompanying transcriptomic changes that occur in different skeletal cell types. We show that although Dlx family factors are playing important roles in classical osteoblast regulation, Hox family factors are involved in governing spinal ossification via non-classical osteoblasts. We further present a resource-driven analysis of the *entpd5a* promoter, experimentally validating the ATAC-seq dataset and proposing mechanisms of regulating the complex *entpd5a* expression pattern during zebrafish osteogenesis. Our results thus provide a necessary comprehensive resource for the field of bone development and indicate spatio-temporally regulated promoter/enhancer interactions taking place in the *entpd5a* locus.

## Introduction

Osteogenesis is one of the vital processes taking place in vertebrate embryos. In amniotes, including all mammals, osteoblasts are known to give rise to all cranial and axial bones through endochondral and intramembranous ossification (Mackie et al., 2008, Breeland et al., 2022). While most cranial bones form via intramembranous ossification, mammalian axial bones are formed via a cartilagenous template by osteoblasts derived from the somitic sclerotome. Despite the differences in modes of ossification, all osteoblasts are considered as a uniform cell type, experiencing similar ontogenetic stages which are controlled by similar genetic mechanisms (Amarasekara et al., 2021; Hartmann, 2009; Jensen et al., 2010).

Teleost and amphibian processes of endochondral and intramembranous ossification are conserved with amniotes (Weigele and Franz-Odendaal, 2016). Although the osteoblasts, which generate bone in amniotes and anamniotes are considered homologous, relatively recent work in teleost fish identified a bone producing cell type, which has not been described in anamniotes: the notochord sheath cell (NSC) (Grotmol et al., 2005; Lleras Forero et al., 2018). NSCs, also referred to as chordoblasts, form an epithelial layer around the vacuolated cells of the embryonic notochord (Fig. 1A), and secrete the molecules making up the acellular, thick notochord sheath layer. NSCs are directly responsible for forming ossified chordacentra, ossified rings that are positioned in a segmental array within the sheath of the notochord. Chordacentra formation precedes the accumulation of sclerotomal osteoblasts around the notochord, which complete formation of the vertebral centra and in a later step also give rise to the dorsal and ventral arches (Fleming et al., 2015; Grotmol et al., 2005; Lleras Forero et al., 2018). Thus, formation of chordacentra is now considered the first stage of spine formation in teleost fish.

**Figure 1.**
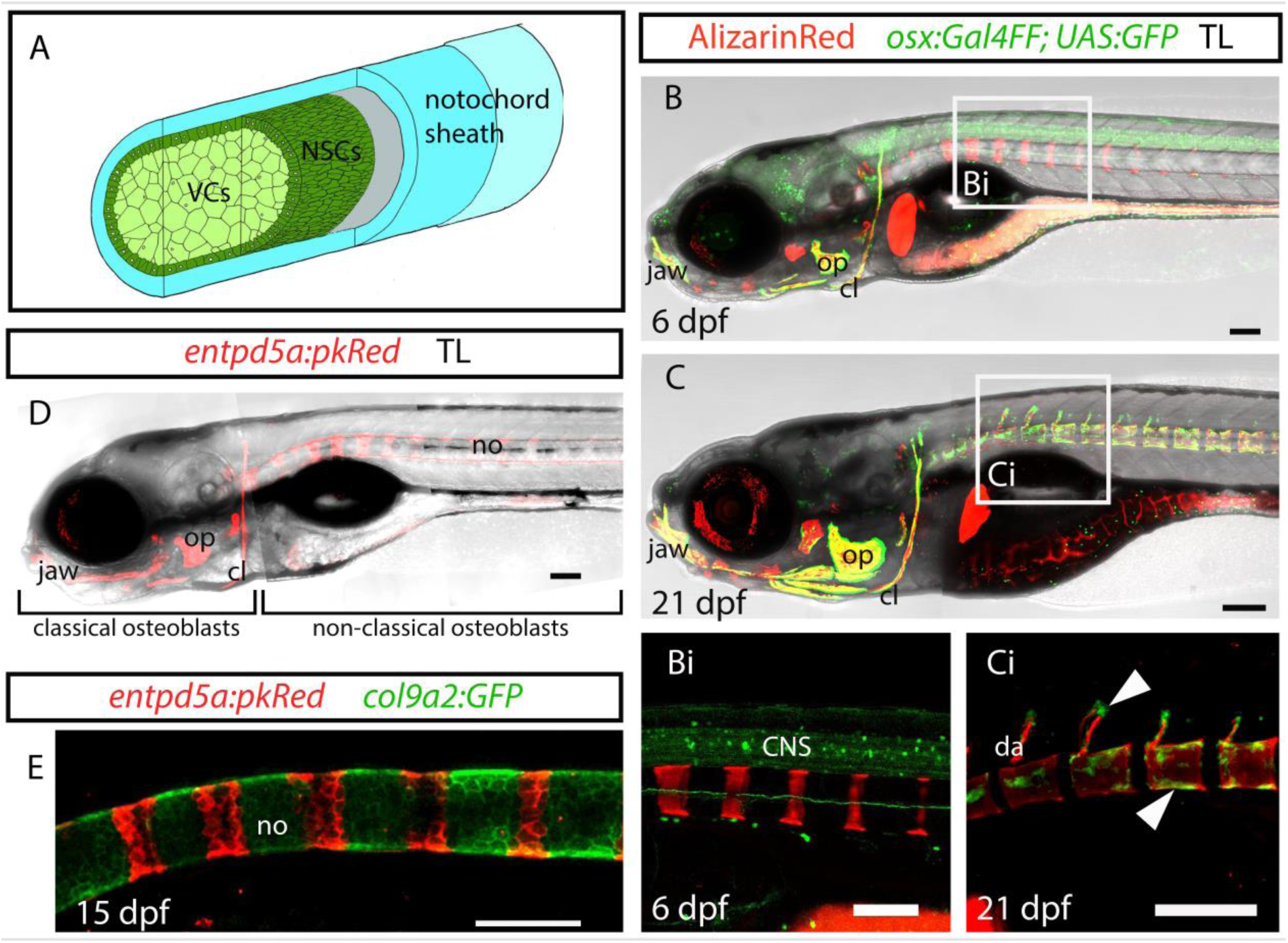
*entpd5a* but not *osterix* is a marker for a non-classical osteoblast population in zebrafish. (A) Schematic of the embryonic notochord structure (adapted from Grotmol et al., 2005). The “notochord sheath” comprise the three outer layers, specifically the basal lamina (grey), the medial layer of fibrillar collagen (light blue) and the outer layer of loosely arranged matrix (lighter blue). Alizarin red staining of mineralised bone reveals non-classical osteoblasts along the trunk that are negative for *osterix* (B, Bi) and positive for *entpd5a* (D) at 6 dpf, as well as classical osteoblasts of the head and mature vertebrae (arrowheads) that are positive for *osterix* (C, Ci) (21 dpf). (E) *entpd5a* and *col9a2* label bone-producing and intersegmental NSCs, respectively. cl, cleithrum; CNS, central nervous system; da, dorsal arch; no, notochord; NSCs, notochord sheath cells; op, operculum; VCs, vacuolated cells. All lateral views, anterior to the left. (B, Bi, D, E) scale bars: 100μm. (C, Ci) scale bars: 200μm.

Here we coin the terms “classical” and “non-classical” osteoblast to describe zebrafish bone formation. Classical osteoblasts, highly homologous to the mammalian osteoblast, express *osterix* and generate the cranial bones and the late perichordal centra of the spine, while non-classical osteoblasts, with to date no known homologous cell type in amniotes, refer to the *osterix-* subset of NSCs capable of early chordacentra mineralisation. Zebrafish cranial classical osteoblasts originate from the cranial neural crest, share a common progenitor with cranial chondrocytes, and can be detected as early as 2 dpf using markers such as *runx2b*, *entpd5a* and *osterix* (Huitema et al., 2012; Li et al., 2009). Axial classical osteoblasts are derived from the somitic sclerotome and contribute to the mineralization of the finite vertebrae in a process very similar to the one observed in mammals. They are rather late in appearance, and *osterix*+ cells can only be found along the growing vertebral central and arches from approximately 17 dpf onwards (Spoorendonk et al., 2008) (Fig. 1B, C).

Non-classical osteoblasts are derived from notochord progenitor cells (NPCs). Unlike classical osteoblasts, NSCs are distinct in their lack of expression of the transcription factor Osterix, commonly considered as an osteoblast master regulator in vertebrates (Nakashima et al., 2002; Wopat et al., 2018). Rather than *osterix*, expression of the early zebrafish bone marker *ectonucleoside triphosphate diphosphohydrolase 5a (entpd5a)* was demonstrated in bone-forming NSCs using BAC transgenic lines (Fig. 1D). As early as 4 dpf *entpd5a* was shown to be expressed in NSCs in a segmental pattern, gradually progressing from anterior to posterior, predicting the precise location of the mineralised chordacentra (Lleras Forero et al., 2018; Wopat et al., 2018). *entpd5a*+ segments along the notochord are separated by intersegmental regions expressing cartilage markers, such as *col9a2* (Fig. 1E). *entpd5a* codes for a secreted hydrolase binding nucleoside di- and triphosphates, generating NMP and inorganic phosphate, a key mineral to initiate the mineralisation process of bone. Indeed, *entpd5a* mutant zebrafish embryos present with a complete lack of ossification (Huitema et al., 2012). In mouse mutants this phenotype has not been reported, although it was observed that homozygous mutants are smaller than their littermates (Read et al., 2009). Thus, NSCs, despite providing a morphologically uniform population of cells, are critical for initial chordacentrum ossification by expressing *entpd5a* in a segmental manner along the anterio-posterior axis. How this segmental expression pattern of *entpd5a* is established is not understood, but it is distinct from the one involved in somitogenesis (Lleras Forero et al., 2018). The role of the BMP and Notch signalling pathways has been highlighted in the segmenting notochord (Peskin et al., 2023; Wopat et al., 2018), but the transcriptional events governing *entpd5a* expression in classical osteoblasts and in the NSCs remain unknown.

The transcriptomics of mammalian classical osteoblasts have been investigated in some detail in the past decades. Genes such as Runx2, Dlx family factors and Osterix play significant roles in different stages of osteoblast specification and maturation (Hassan et al., 2004; Hojo et al., 2022; Jensen et al., 2010; Li et al., 2008; Shirakabe et al., 2001). More recently, the Assay for Transposase-Accessible Chromatin (ATAC-seq) has been widely used in the field. In addition, an increasing number of studies integrate RNA-seq and ATAC-seq to provide deeper insight into gene regulation in populations of interest (Ackermann et al., 2016; Hao et al., 2022; Hojo et al., 2022; Yang et al., 2021; Yu et al., 2021). Such studies, however, are to date available neither for classical, nor for non-classical zebrafish osteoblasts, rendering the molecular fingerprints and chromatin dynamics of zebrafish osteogenic cells to date elusive. Molecularly characterising zebrafish classical osteoblasts (Raman et al., 2024) is of particular interest for the field, as it allows for direct comparison between mammalian and teleost transcriptional regulation of osteogenesis. Specifically, it is crucial to acquire a deeper understanding of teleost osteoblasts in order to answer evolutionary questions regarding the cell type’s origin (Nguyen and Eames, 2020). Moreover, better understanding the mechanisms of regulation in non-classical osteoblasts is likely to highlight either novel ossification mechanisms that have not been appreciated to date, or that are unique in teleosts.

In this study, we provide the first comprehensive resource of ATAC-seq and RNA-seq performed on classical and non-classical zebrafish osteoblasts, enabling studies of embryonic osteoblast development and differentiation. Comparisons across those populations and their closely related chondrocyte and intersegmental NSC populations allows a detailed assessment of gene expression profiles and chromatin regulation in different osteogenic cells, based on FAC-sorted cells from live embryos. We furthermore focus on the promoter structure of the early bone marker *entpd5a*, a gene critical for osteogenesis in zebrafish. We thus not only validate the usefulness and quality of the ATAC-seq data, but also demonstrate the complex mode of *entpd5a* regulation through identification of enhancers located both upstream of the ATG and within intronic regions, and suggest candidate transcription factors likely to be acting on the identified enhancer elements.

## Results

### *entpd5a* is an early marker of developing osteoblasts

Entpd5a is a critical factor for zebrafish osteogenesis, and *entpd5a* mutants show no bone formation at all. We had previously generated a transgenic reporter line and established the gene to be expressed at sites of bone formation, but wanted to better understand the dynamics of *entpd5a* expression. We thus recombined the BAC construct CH73-213B8, which was used previously to generate stable *entpd5a* transgenic lines (Bussmann and Schulte-Merker, 2011), and replaced the start codon of the gene with that of the Gal4FF open reading frame (Suppl. Fig. 1A). The Gal4/UAS system has proven valuable for other genes to provide stronger fluorescence signals. Indeed, the newly generated *entpd5a:Gal4FF*; *UAS:GFP* transgenic line (Labbaf et al., 2022) allowed detection of *entpd5a* expression domains which had previously not been reported.

Using this line, we could detect for the first time expression of *entpd5a* not only in the developing cranial bones starting from 3 dpf, but also in a subset of cartilage cells forming the developing jaw (Suppl. Fig. 1B, D). This pattern is reminiscent of the one previously shown for the bone master regulator *osterix* (Hammond and Schulte-Merker, 2009). In addition, we detected *entpd5a* expression in a few neurons of the CNS in the head at 3 dpf (Suppl. Fig. 1C) and along the spinal cord. We also found expression within the hypochord (data not shown). Finally, observation of the strongly expressing Gal4 line led us to identify early expression of *entpd5a* in notochord progenitor cells (NPCs) at 24 hpf (data not shown).

To further elucidate early *entpd5a* expression, we used the previously established *entpd5a:Kaede* transgenic line (Lleras Forero et al., 2018) to investigate notochord dynamics prior to 3 dpf. We photoconverted and directly imaged *entpd5a:Kaede* positive embryos starting from the 15 somite-stage (s), when we could first detect the fluorophore along the newly-formed notochord progenitor cells (Suppl. Fig. 1E). We repeated photoconversion and imaging at 18, 21 and 24s (Suppl. Fig. 1F-H). In all three stages, *de novo* expression (in green) was identified. We repeated the photoconversion and imaging at 48 hpf and at 72 hpf (Suppl. Fig. 1I-J). At both stages, expression of *entpd5a* was restricted exclusively to NSCs positioned along the ventral-most notochord sheath. Hence, the previously observed expression of *entpd5a* in the notochord prior to segmentation (Lleras Forero et al., 2018) in vacuolated cells and a subset of NSCs is merely the result of long half-lives of fluorophores.

In conclusion, *entpd5a* expression in the zebrafish embryo appears to be more dynamic than previously appreciated, which implies diverse regulatory mechanisms. A list of tissues expressing *entpd5a* (Table 1) provides a better understanding of *entpd5a* expression dynamics, and serves as a basis for considering ATAC- and RNA-sequencing datasets.

**Table 1.**
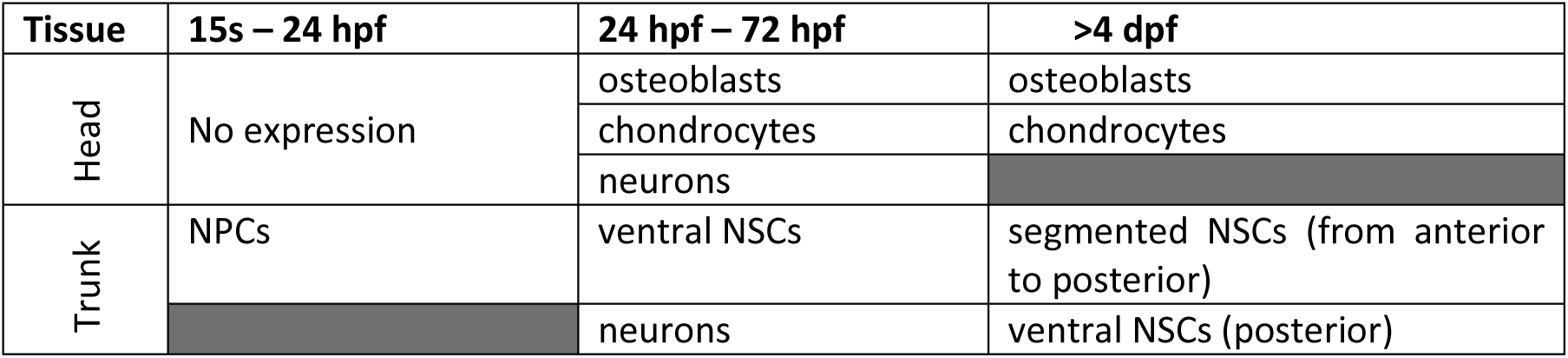
Revised list of tissues with detected *entpd5a* in the developing zebrafish embryo.

### ATAC-seq on classical versus non-classical osteoblasts identifies a high degree of correlation between regions of open chromatin

In order to identify cell type-specific enhancers in the different cell types forming the zebrafish skeleton, and to further predict crucial transcriptional regulators that are active in each population, we performed ATAC-seq using cells isolated from the *entpd5a*:*Gal4FF; UAS:GFP*; *R2col2a1a:mCherry* transgenic line at 15 dpf (Suppl. Fig. 2A-B). The developmental stage was chosen in order to obtain sufficient numbers of *entpd5a+* and *entpd5a-* NSCs (Suppl. Fig. 2A). Larvae were decapitated and dissociated, and cells were sorted according to fluorescence, using GFP-; mCherry- embryos as controls. Classical GFP+ osteoblasts and mCherry+ chondrocytes were isolated from the head and GFP+ NSCs and mCherry+ intersegmental NSCs from the trunk (Suppl. Fig. 3).

We assessed the quality of sequenced data according to ENCODE standards and used ATAQV (Orchard et al., 2020) to determine the Fragment Length Distribution Distance (FLDD) and the TSS enrichment score (Supplementary Table 1). The FLDD scores were always found to be between 0 and 0.3, indicating good degree of transposition. The TSS value, although low in comparison to data from mammalian cells, indicates enrichment of fragments around the promoters. Furthermore, we assessed sample correlation via Principal Component Analysis (PCA) and by producing a correlation heatmap based on FRiP (Fraction of Reads in Peaks) scores using DiffBind (Suppl. Fig. 2C-D). Both methods highlighted a high degree of correlation between biological replicates, as well as tight clustering between all 6 samples derived from *entpd5a*+ cells versus the 6 samples derived from *entpd5a*- cells. These correlations confirmed the reliability of the data.

Differential accessibility analysis was performed, comparing the two head-derived cell populations with each other to identify putative enhancer regions associated with either classical osteoblasts or with chondrocytes (Suppl. Fig. 2E). Similarly, the two trunk populations were compared to identify open chromatin regions specific to the *entpd5a+* bone-producing NSCs, and chromatin regions specific to *entpd5a-* intersegmental NSCs (Suppl. Fig. 2F). Overall, we identified here 558 peaks specific to classical osteoblasts and 26 peaks enriched in NSCs (non-classical osteoblasts). The latter small number is likely due to the high degree of similarity of the two cell types compared (*entpd5a*+ vs. *entpd5a*- NSCs).

### Functional validation of ATAC-seq leads to novel insights into *entpd5a* regulation

To test if the ATAC-seq open chromatin regions are predictors of putative regulatory elements, we focused on the *entpd5a* locus, and used GFP reporters to perform enhancer function tests *in vivo*. Within close proximity to the *entpd5a* ORF six ATAC peaks show enrichment in osteoblasts and represent enhancer candidates (Fig. 2A). Further away from the ORF, the non-coding sequences present in the BAC contain an additional 2 peaks of interest for osteoblast regulation (peaks 7-10, Fig. 2B) and 2 peaks enriched in cartilage samples (peaks 8/9, Fig. 2B). Using the ZF_carp_phastcons track on the UCSC browser (Baranasic et al., 2022), we assessed the degree of teleost non-coding sequence conservation in close proximity to the open reading frame of *entpd5a*. We found that all 10 ATAC-peaks were positioned within conserved non-coding regions, suggesting the presence of cis regulatory elements (Fig. 2B). We first focused on ATAC peaks 1-6 within approximately 10kb of the *entpd5a* ATG (Fig. 2A). As shown in the diagram, the first peak was found in an intron at the 3’ end of the *coq6* gene. Peaks 2 and 3 were contained in introns 3 and 1, respectively, of the *entpd5a* gene. The first introns of a gene often play regulatory roles (Rose, 2019), making these peaks intriguing. Finally, we highlight peak 4, close to the TSS of *entpd5a*, (Nepal et al., 2013) as well as two more peaks (5 and 6) located within 6kb and 11kb of the TSS, respectively. We sought to experimentally test possible roles of these open chromatin domains, by generating transgenes that place these domains 5’ or 3’ of a GFP casette, and comparing the resulting expression patterns upon reporter transgenesis to that of the *entpd5a* locus (Table 1).

**Figure 2.**
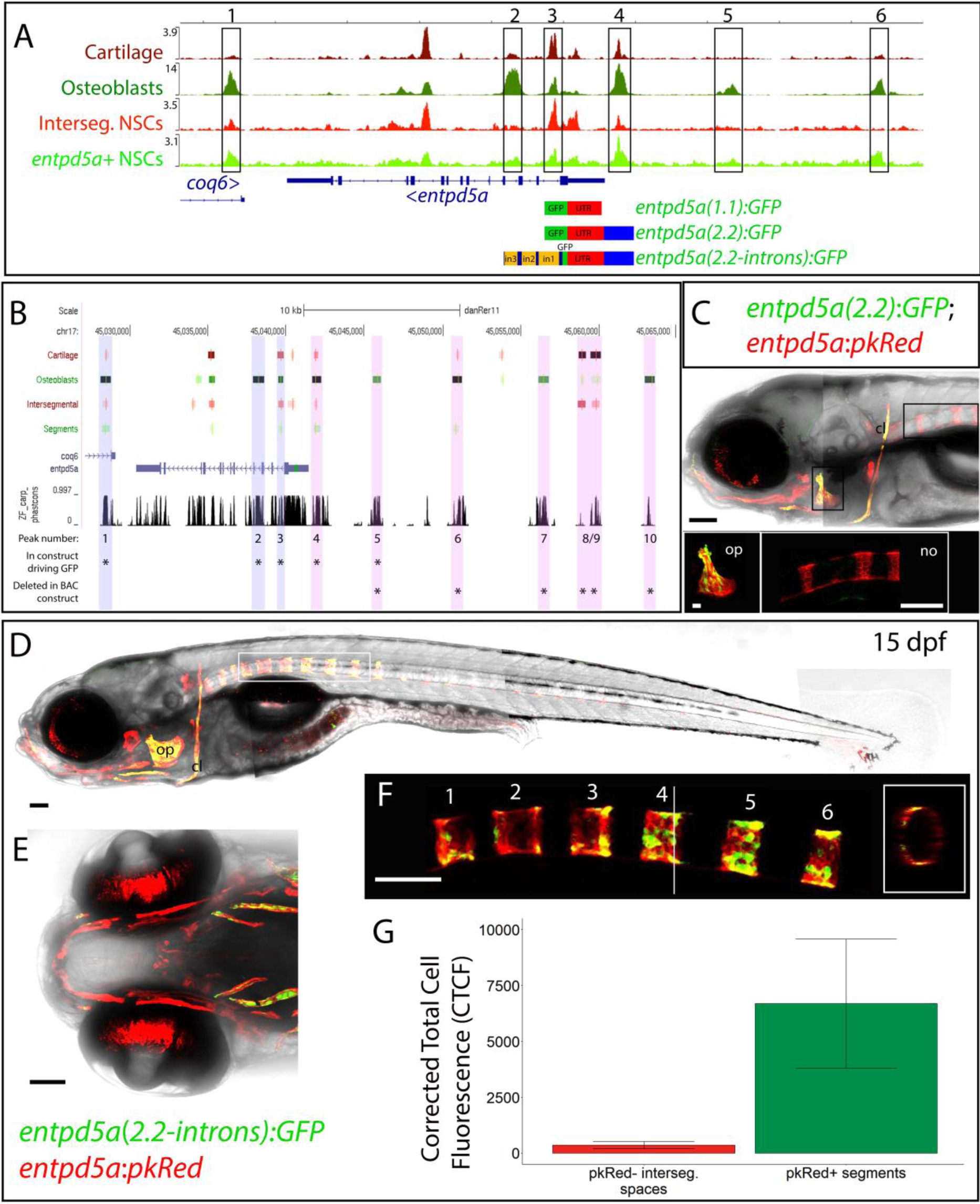
Validation of potential enhancer regions in the *entpd5a* promoter indicates distinct regulation of classical and non-classical osteoblasts. (A) Chromatin accessibility profiles of representative samples of each cell type. Highlighted are six differentially accessible regions between *entpd5a*+ and *entpd5a*-cells. Chromatin regions 4-6 lie proximal to the *entpd5a* ORF, regions 2-3 are positioned within introns 1 and 3 respectively, whereas region 1 is found in the 3‘ end of the *coq6* gene. The three construct schematics (*entpd5a(1.1):GFP, entpd5a(2.2):GFP, entpd5a(2.2- introns):GFP*), indicate the sequences cloned for generation of the corresponding transgenic lines.(B) View of the UCSC browser showing a total of 10 regions of open chromatin within the 40kb surrounding the ORF of *entpd5a,* including regions 1-6 shown in (A) and regions 7-10 located further upstream. ORFs of entpd5a and downstream *coq6* are shown in purple. The TSS of *entpd5a* is marked as a green bar in the 5’UTR. The ZF_carp_phastcons track indicates conservation amongst zebrafish, goldfish and 2 species of carp. Peaks identified by *macs2* as significant compared to background are shown as boxes for each sample (red tracks for cartilage and intersegmental cells and green tracks for osteoblasts and segments). Boxes darker in colour indicate peaks with higher score values. Peaks of interest 1-10 are highlighted and it is indicated using asterisks whether a peak was tested for enhancer function by inserting the sequence in a construct driving GFP and/or by deletion in the BAC construct. (C) Embryo from a stable transgenic line with region 4 placed upstream of GFP, imaged at 6 dpf. GFP is expressed in osteoblasts (marked using *entpd5a*:*pkRed*), but not in the notochord. Insets demonstrate GFP and pkRed overlap in the operculum (op), but not in pkRed+ segmented NSCs. (D, F) lateral and (E) ventral head view of stable line where GFP is driven both by region 4 and by regions 2 and 3. Cranial osteoblasts are expressing GFP (D, E), and segmentation is clearly observed in the notochord (D, F). (G) Quantification of fluorescence in segments 1-6 versus intersegmental regions as shown in (F). p-value = 0.003. cl, cleithrum; no, notochord; op, operculum. (C, D, F) lateral views; (E) ventral view. Anterior to the left in all images. (C-F) scale bars: 100μm. Operculum (C) inset scale bar: 20μm.

We first generated an *entpd5a(2.2):GFP* construct by cloning a 2.2kb fragment including the 5’UTR and core promoter of *entpd5a* as well as the non-coding region upstream of it, which includes peak number 4. The fragment was placed upstream of the GFP open reading frame (Fig. 2A), with Tol2 sites on either side of the construct. As a control, we generated *entpd5a(1.1):GFP*, a construct containing only the 5’UTR, along with the TSS and core promoter of the gene. We generated stable transgenic lines by injecting *entpd5a(1.1):GFP* or *entpd5a(2.2):GFP* together with Tol2 transposase mRNA into zebrafish eggs. In fish stably expressing *entpd5a(1.1):GFP*, expression could be detected in a variety of tissues but not in regions of ossification (data not shown). The result was confirmed by analysing progeny of six separate founders. In *entpd5a(2.2):GFP; entpd5a:pkRed* stable lines, we identified overlap of GFP and pkRed in the developing cranial bones, specifically the cleithrum, the operculum and a subset of jaw bones, but not in the notochord (Fig. 2C). While at 6 dpf segmentation in the notochord is ongoing, as can be readily observed using the *entpd5a*:*pkRed* stable line, GFP is undetectable in pkRed+ NSCs (Fig. 2C, inset). This result was consistently found in progeny of three different founders, indicating the presence of proximal enhancers of *entpd5a* functioning in cranial osteoblasts in the sequence where peak number 4 is identified.

In order to assess whether additional *entpd5a* enhancers are present upstream of the area covered by the *entpd5a(2.2):GFP* construct, we generated *entpd5a(5.7):GFP*, which extends a further 3.5kb, covering the peak number 5 sequence (Suppl. Fig. 4A). The sequence covered by the *entpd5a(5.7):GFP* construct is conserved across teleosts, but conservation is no longer observed 5’ of the cloned fragment (Fig. 2B). We generated stable transgenic lines, and imaged at 6 dpf in the background of the *entpd5a:pkRed* line, to identify cranial expression, as well as notochord expression (Suppl. Fig. 4B, C). As for *entpd5a(2.2):GFP*, we could only identify GFP expression in pkRed+ developing cranial bones, but not in the notochord (Suppl. Fig. 4B, C). Furthermore, although pkRed was detectable in the jaw chondrocytes, GFP signal was not found, indicating lack of a cartilage-specific enhancer capable of initiating expression autonomously within the sequence of peak number 5 (Suppl. Fig. 4B, inset). These results were validated by assessment of the progeny of eight founders. In conclusion, conserved sequences amongst teleosts upstream of peak number 4 do not appear to contain any enhancers that play roles in activating expression of *entpd5a* during development.

We next tested genomic regions equivalent to peaks 2 and 3, found in teleost-conserved *entpd5a* introns 3 and 1, respectively, for activating regulatory functions. To this end, we amplified the region between exon 1 and exon 4, including intron 1-3, and introduced them in the *entpd5a(2.2):GFP* construct, directly downstream of the GFP open reading frame, thus generating the construct *entpd5a(2.2-introns):GFP*. We injected the construct into single cell-stage zebrafish eggs, and observed that NSCs were expressing GFP in a mosaic manner (Fig. 3F-G), an effect not seen with other constructs. We generated an *entpd5a(2.2-introns):GFP* stable transgenic line, and indeed the notochord expression of *entpd5a*, as observed in the *entpd5a*:*pkRed* transgenic line, was present from 24 hpf until segmentation stages (Suppl. Fig. 5, Fig. 2D-F). Although GFP appeared in the expected segmental pattern along the notochord, expression levels were relatively low and patchy. This may be the result of this specific integration event (only one founder could be identified), or due to lack of additional enhancers located in other non-coding regions upstream or downstream of the gene. We confirmed, however, that the expression of GFP was enriched in segments compared to intersegmental spaces, by measuring the fluorescence intensity within 6 segments and 5 intersegmental spaces, respectively (Fig. 2F). Plotting the corrected total cell fluorescence measurements in each region (Fig. 2G) confirmed the segmental pattern of GFP in Fig. 2F.

**Figure 3.**
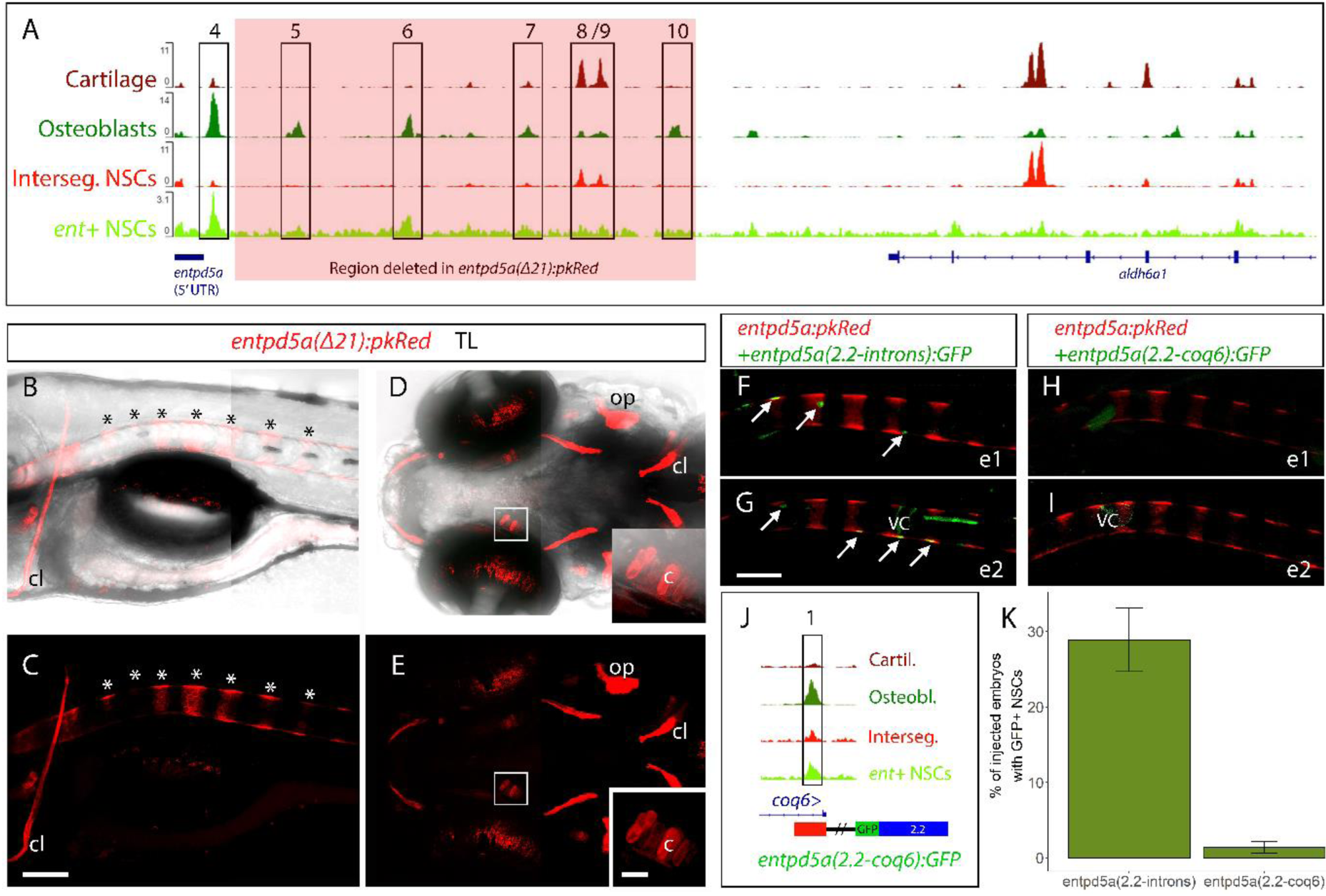
Deletion of open chromatin further upstream or downstream of the *entpd5a(2.2-introns)* domain does not affect the *entpd5a* expression pattern. (A) Chromatin accessibility profiles along the length of the *entpd5a* BAC, starting from the ATG of *entpd5a*. Representative tracks for each cell population are shown. Peaks of interest are labelled 4-10, following the labelling in Fig. 2. The region highlighted in red indicates the 21.4kb deleted in the BAC construct *entpd5a(Δ21):pkRed*. (B-E) Embryo stably expressing the *entpd5a(Δ21):pkRed* construct shows normal segmentation pattern at 6 dpf (B-C, asterisks), as well as normal osteoblast and chondrocyte (D, E, insets) expression of pkRed. (F-H) two representative *entpd5a:pkRed+* embryos injected with *entpd5a*(*2.2-introns):GFP* show mosaic GFP expression along NSCs associated with pkRed+ segments at 5 dpf (F, G). In contrast, representative embryos injected with *entpd5a*(*2.2-coq6):GFP* only show background GFP at the same stage (H, I). (J) Schematic of the *entpd5a*(*2.2-coq6):GFP* construct, integrating peak number 1 downstream of the GFP. (K) Proportion of embryos injected with GFP constructs with observable GFP expression in NSCs. c, chrondrocyte; cl, cleithrum; op, operculum; VC, vacuolated cell. (B, C, F-I) lateral views; (D, E) ventral views. Heads positioned towards the left. (B-I) scale bars 100μm. (D, E) insets’ scale bar: 20μm.

Neither of the above constructs showed GFP expression in jaw cartilages, thus, a cartilage-specific enhancer of *entpd5a* has not been identified. We performed BAC recombineering to delete larger regions of the *entpd5a*:*pkRed* BAC, in order to assess presence of enhancers within peaks 6-10, as well as within the peak found in intron 11, which appears more prominent in cartilage cell populations (Fig. 2B, 3A). First, we generated the large deletion *entpd5a(Δ21):pkRed*, spanning 21.4kb upstream of the *entpd5a(2.2):GFP* construct, deleting the sequence harbouring peaks 5-10 from the BAC construct (Fig. 3A). We generated a stable transgenic line and found that the expression pattern appeared unaltered, with strong expression appearing in the cranial skeleton, cartilage and notochord (Fig. 3B-E). Therefore, no additional upstream open chromatin regions were found to contribute to the wild type expression pattern as depicted in Table 1. Moreover, we generated the deletion *entpd5a(Δ10):pkRed,* in which the intron 11 sequence was removed, but this failed to affect expression of pkRed in cartilages (data not shown).

Moreover, we went on to assess in a comparable *in vivo* setting the presence of active *entpd5a* enhancers within the region of peak number 1, located in the last intron of the gene *coq6* (Fig. 2A). Similarly to the generation of *entpd5a(2.2-introns):GFP*, we generated the construct *entpd5a(2.2- coq6):GFP* (Fig. 3J), and injected it in single cell-stage zebrafish embryos. We scored the fish injected with either of the two constructs and found that NSCs were not expressing GFP in the case of *entpd5a(2.2-coq6):GFP*, while such cells were commonly occurring when the *entpd5a(2.2-introns):GFP* was injected (Fig. 3F-I, K).

We thus conclude that out of the 10 peaks of interest (Fig. 2B), only ATAC peaks 2, 3 and 4 (Fig. 2A) identify genomic elements playing activating roles in *entpd5a* regulation, with proximal enhancers in peak number 4 driving classical and regulatory elements in peaks 2 and 3 non-classical osteoblast expression. We further found that the open chromatin located further 5’ of the *entpd5a* TSS or downstream of the open reading frame plays no role in regulating expression in the developing cartilage and bone tissue of zebrafish embryos.

### Identification of the classical osteoblast proximal enhancer region

In order to further narrow down the region within the *entpd5a(2.2):GFP* construct, which is responsible for driving cranial expression of *entpd5a*, we generated sequential deletions from the 3’ end of the construct. Constructs *entpd5a(d243):GFP*, *entpd5a(d418):GFP*, *entpd5a(d665):GFP* and *entpd5a(d880):GFP* carry deletions of 243bp, 418bp, 665bp and 880bp, respectively (Fig. 4A). Using the CIIIDER program (Gearing et al., 2019), which predicts TF binding sites along a sequence of interest, we identified two elements along the 2.2kb sequence that were predicted to contain putative binding sites of Runx2, a known regulator of osteogenesis. We wished to test whether these sites were functional for cranial osteoblast expression of *entpd5a* and generated three additional constructs, *entpd5a*(*r37):GFP*, *entpd5a(r31):GFP* and *entpd5a(r37/r31):GFP*, respectively carrying a 37bp deletion at the first Runx2 site, a 31bp deletion at the second Runx2 site, and both deletions simultaneously (Fig. 4A).

**Figure 4.**
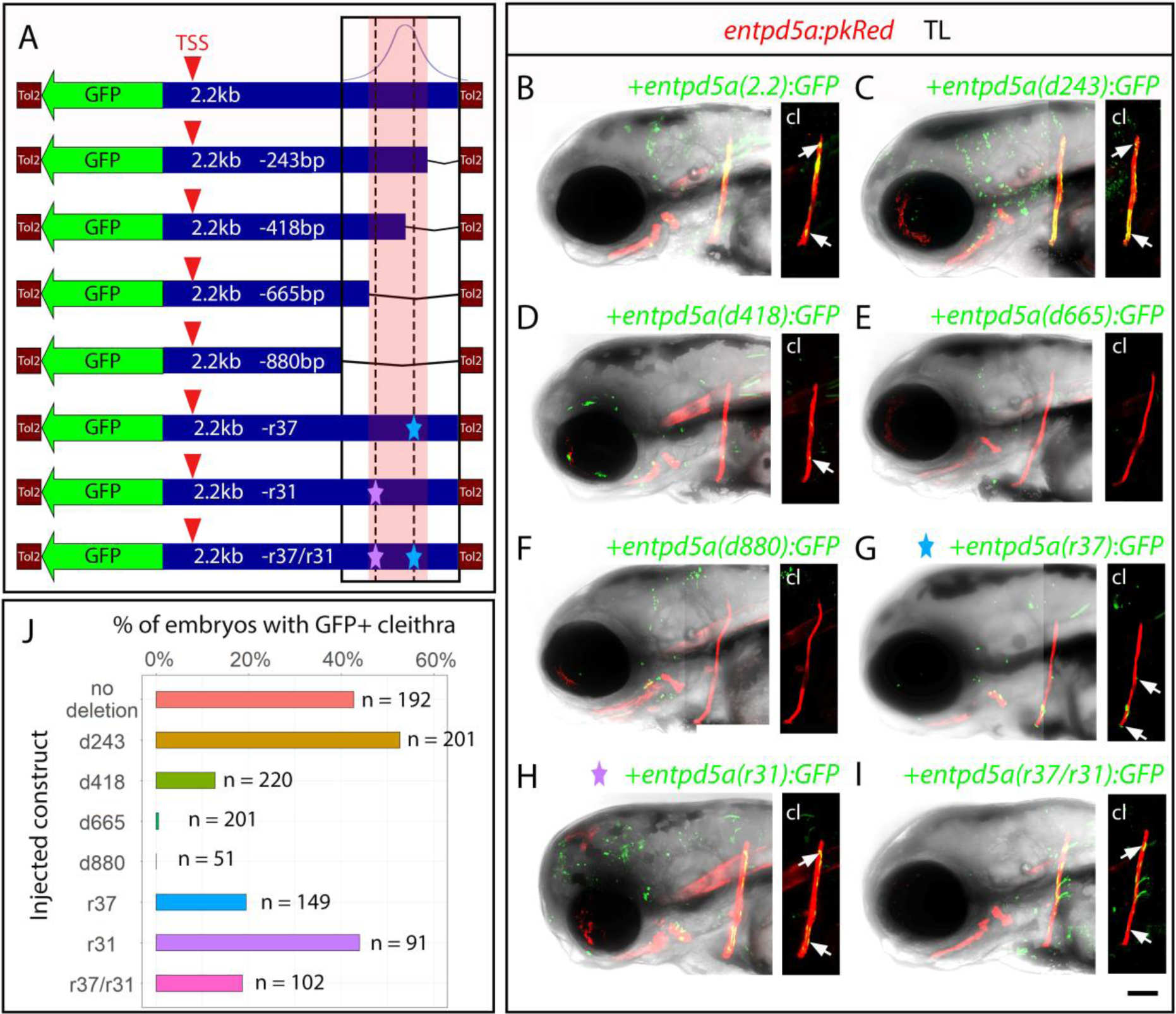
Sequential deletions of the *entpd5a(2.2):GFP* construct highlight a 422bp region containing active *entpd5a* enhancers. (A) Schematic of the initial construct (2.2kb cloned upstream of GFP), and the subsequent deleted constructs. In the final construct, the purple star depicts the -31bp deletion and the light blue star the -37bp deletion. The osteoblast ATAC peak position is indicated. The 422bp of interest is highlighted in red. (B-I) Representative *entpd5a:pkRed* embryos at 3 dpf injected with the respective deletion constructs, or as a control with the complete construct (B). The cleithra (cl) for each embryo are depicted without transmitted light. (J) Graph indicating, across all experimental repeats, the percentage of successfully injected embryos (n) for each injected construct in which GFP+ cells were observed in the cleithrum. Successfully injected embryos were identified based on the presence of GFP+ cells in background tissues. Lateral views, anterior to the left. (B-I) scale bar: 100μm.

We injected the constructs into zebrafish embryos carrying *entpd5a*:*pkRed* at the single cell-stage, and screened at 3 dpf for GFP expression in the cleithrum, a morphologically distinct bone of the zebrafish skeleton. The total numbers of screened embryos with GFP+ cells along their bodies, indicating successful integration of the construct, are provided in Fig. 4J, and are derived from a minimum of 2 biological replicates for each construct. Each injection was accompanied by independent injections of the *entpd5a(2.2):GFP* control construct in sibling embryos. Of the GFP-expressing embryos, we scored embryos as positive when 2 or more GFP+ cells were present along the cleithrum at 3 dpf. When the *entpd5a(2.2):GFP* and the *entpd5a(d243):GFP* constructs were injected, 42.7% and 52.7% of the scored fish presented with strongly GFP-positive cleithra (Fig. 4B, C, J). The number of scored injected fish with GFP in their cleithra dropped to 12.7%, 0.5% and 0% when constructs *entpd5a(d418):GFP*, *entpd5a(d665):GFP* and *entpd5a(d880):GFP*, respectively, were injected (Fig. 4J). We observed that in those injections only very few GFP+ cells (if at all) were detected along the cleithra, in stark contrast to the control and *entpd5a(d243):GFP* constructs (Fig. 4B-F). This indicates that a region of 422bp (highlighted as a red box in Fig. 4A) is responsible for *entpd5a* regulation in classical osteoblasts.

The identified 422bp region of interest includes two putative Runx2 binding sites. We injected the three constructs, deleting them both individually and simultaneously, and found that, although deletion r31 had no effect in the number of scored embryos with GFP expression in the cleithrum at 3 dpf (44%), deletions r37 and r31/r37 both resulted in a consistent drop to approximately half of the control group (19.5% and 18.6%, respectively) (Fig. 4J). This led us to conclude that the predicted Runx2 binding site located close to the summit of the ATAC peak has a functional role in *entpd5a* expression in the cranial skeleton.

### Integration of ATAC-seq and RNA-seq identifies candidate regulators of classical and non-classical osteoblasts

In order to identify possible transcriptional regulators active in cell types of interest, we sought to integrate our ATAC-seq dataset with differential expression data from RNA-seq. To that end, we performed RNA-seq at 15 dpf on cranial osteoblasts and cartilage chondrocytes, as well as on *entpd5a*+ NSCs and intersegmental *entpd5a*- NSCs. As with ATAC-seq, biological triplicates of all samples were sequenced and analysed.

Our RNA-seq results highlighted unique signatures, with *entpd5a*+ cells from head and trunk highly expressing genes known to function in bone formation, for example *phex* (Quarles, 2003), *panx3* (Ishikawa and Yamada, 2017) and *col10a1b*. In contrast, *entpd5a*- cells from head and trunk were found to express genes known to function in cartilage formation, such as *acan* and *ccn* family genes (Dateki, 2017; John A. Arnott, 2016) as well as *col9a1b* (Fig. 5A, B). We further show that, as expected, *entpd5a*+ cells isolated from both tissues share 872 genes, accounting for 43% of the total differentially expressed genes (DEGs) of both head and trunk *entpd5a*+ cells (Fig. 5C). Similarly, *entpd5a*- cells share 765 genes, accounting for 64% of the total DEGs found in cartilage and 40% of those found in intersegmental cells. This high degree of overlap was in contrast to the very low number of genes shared between osteoblasts and intersegmental cells (43 DEGs) or between cartilage and *entpd5a*+ NSCs (7 DEGs). We confirmed the reliability of our results by performing PCA analysis, in which the 3 biological replicates of each cell type clustered together (Fig. 5D). Of note, Principal Component 1 (PC1) distinguishes between the samples isolated from head versus trunk, while PC2 effectively distinguished between *entpd5a*+ and *entpd5a*- cells from both tissues. Overall, we observed significant overlaps in gene expression signatures between the *entpd5a*+ cells and *entpd5a*- cells, even though they are distinctive cell types, originating from widely diverse progenitors. We do note, however, that despite the similarities, classical and non-classical osteoblasts maintain significant differences in their expression profiles.

**Figure 5.**
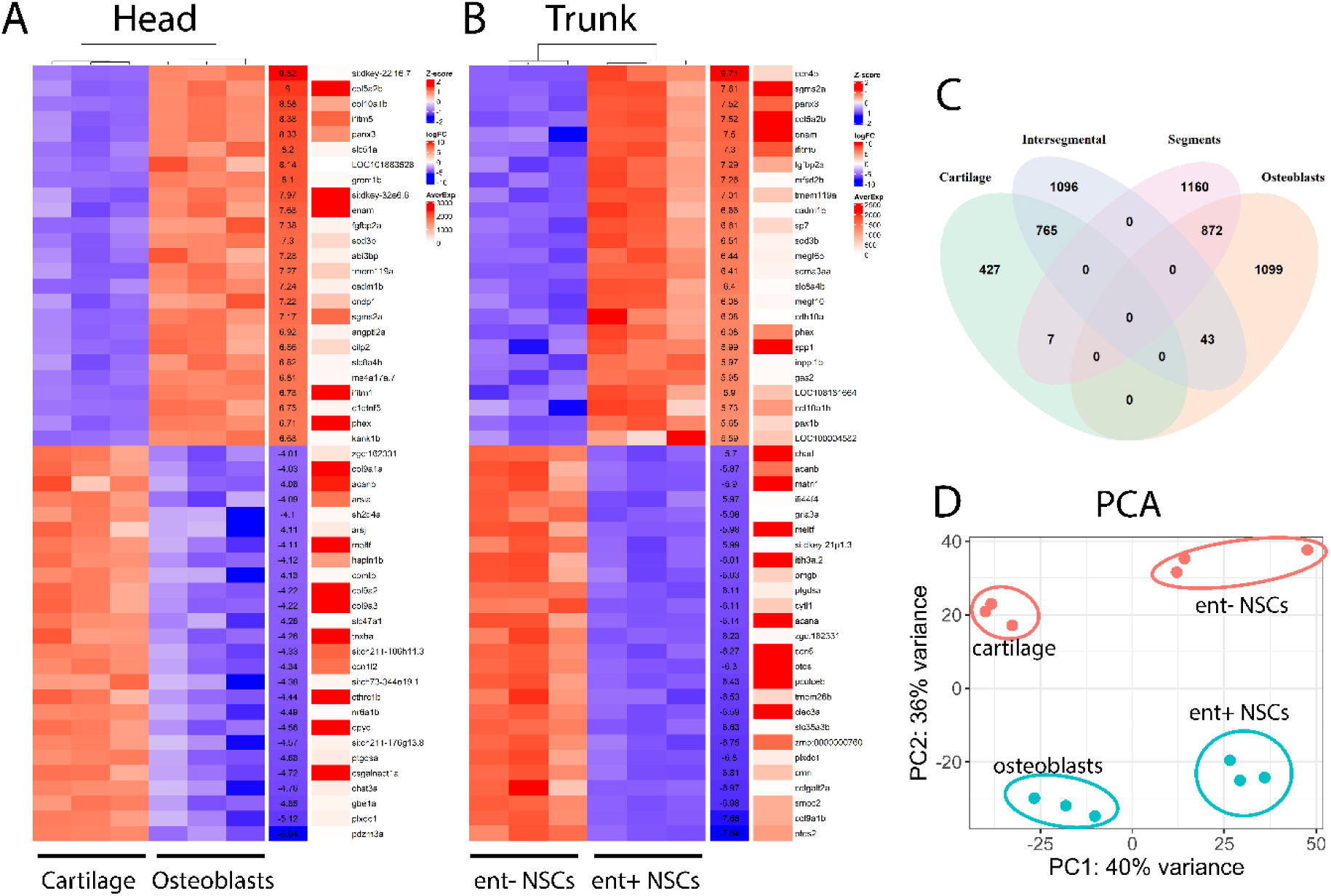
RNA-seq analysis of skeletogenic populations of the head and trunk. (A, B) Heatmaps for each tissue, indicating the top differentially regulated genes. The colours on the main heatmap indicate the Z-score value, while the second heatmap indicates the corresponding logFold change, and the third the Average Expression of each gene. (C) Venn Diagram indicating the overlap of total DEGs in all 4 cell populations. Osteogenic and cartilage tissues share 872 and 765 upregulated genes, respectively, while the non-contrasted osteoblasts vs. intersegmental and cartilage vs. segments share 43 and 7 genes, respectively. (D) PCA plot showing the clustering of cell populations by tissue (head versus trunk) and by *entpd5a* expression (positive and negative cell populations in blue and orange, respectively).

In teleost fish, osteoblasts have been previously found to share expression of typical chondrocyte genes, providing support to theories of bone cells having evolved from cartilage cells (Nguyen and Eames, 2020). Looking closely at our RNA-seq data and ATAC-seq data we found that, despite the logFC values showing clear enrichment in *entpd5a*- cells, indeed classical chondrocyte markers maintain open promoters (Suppl. Fig. 6A-D) and low levels of gene expression relatively to cartilage and intersegmental cells (Suppl. Fig. 6E). It should be noted that ATAC signal indicating the presence of active enhancers in promoter areas is observably lower, often by an order of magnitude, in osteoblasts compared to cartilage cells (Suppl. Fig. 6A-D).

We next combined the information of the ATAC-seq and RNA-seq datasets, to gain further insight into gene regulatory activity in our cell types of interest. To this end, we searched within intronic regions, as well as (a) 5kb, (b) 10kb and (c) 20kb upstream of the TSS of all genes in the GRCz11 genome assembly and isolated the respective three sets of zebrafish genes associated with ATAC peaks. First, we made an assessment of the distribution of peaks in the vicinity of genes. For each cell population, we plotted the proportion of zebrafish genes that had ATAC peaks in close association with promoter regions (within 5kb of the TSS and in the introns), in comparison to the proportion of genes that only had ATAC peaks associated with sequences within 10kb or within 20kb of their TSS (Fig. 6A). We concluded that in all cell populations, the majority of identified genes had peaks close to their promoter regions or within the intronic domains. A relatively small percentage of genes showed peaks only associated with the area 5kb-10kb of the TSS or with the area 10kb-20kb of the TSS. The same conclusion was reached upon isolation from these three isolated sets of genes that are actively expressed in each cell population according to our RNA-seq data. We plotted the proportion of cell type-specific DEGs that are associated with cell type-specific ATAC peaks and found that an even larger proportion (in all populations over 80%) of identified DEGs had ATAC peaks in close proximity to their predicted TSS, compared to very few additional DEGs identified for having peaks in areas between 5kb- 20kb of the TSS (Fig. 6B). This effect further validated our ATAC-seq results, showing that ATAC peaks were concentrated close to the TSS of genes, rather than being distributed evenly in non-coding regions of the genome. Furthermore, we observed that the proportion of genes with peaks within 5kb of their TSS increased when looking exclusively at DEGs, further supporting the higher degree of chromatin accessibility around their promoters, related to their active regulation.

**Figure 6.**
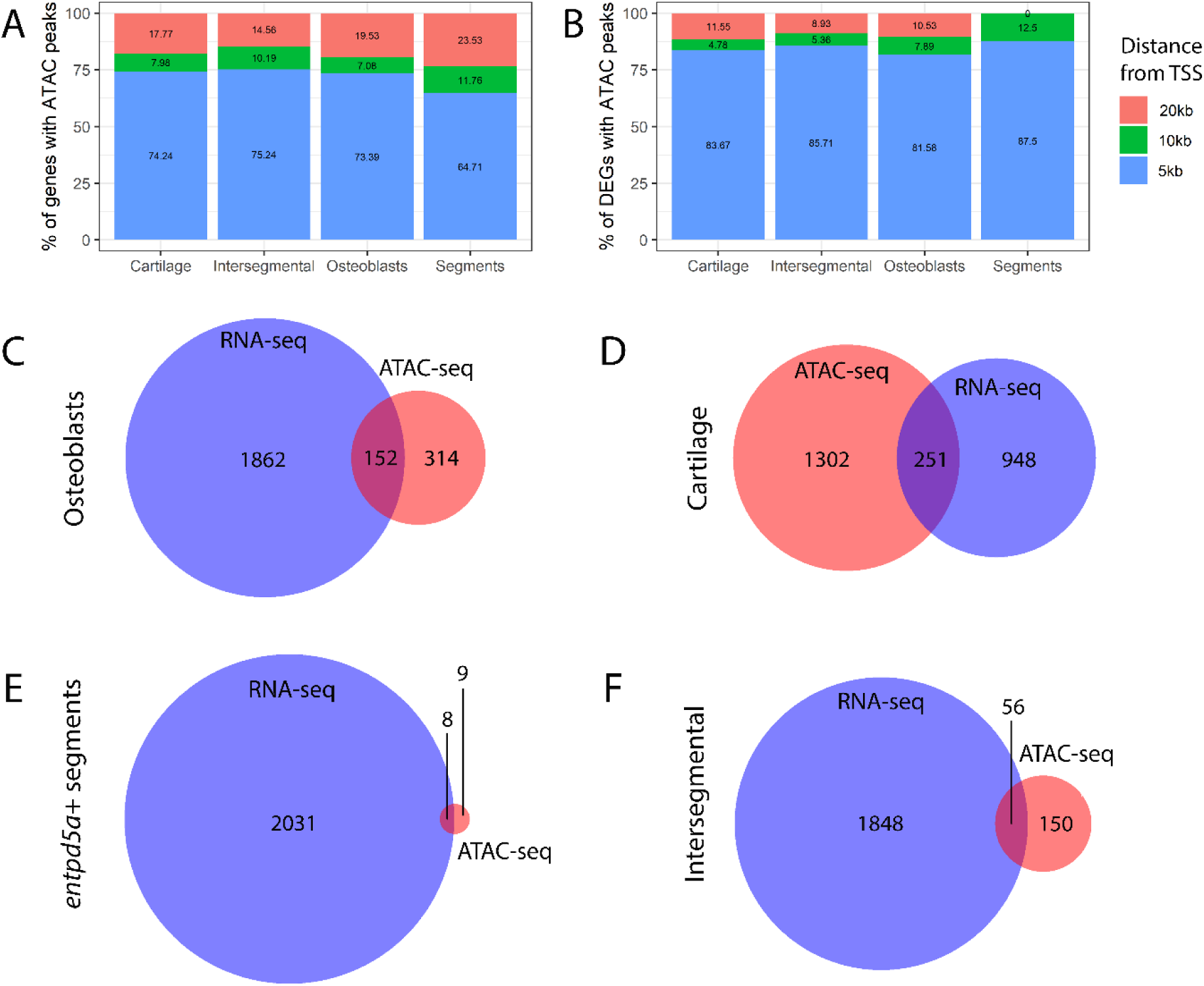
Integration of ATAC-seq and RNA-seq data highlights differentially expressed genes with accessible promoter regions in distinct cell populations. (A, B) Bar graphs indicate the proportion of ATAC peaks found within 5kb, 10kb and 20kb of the TSS in each cell population. In (A) peaks were searched in all of the Zv11 genes and in (B) only in differentially expressed genes (DEGs). (C-E) Venn diagrams indicating for each cell population the overlap between genes in Zv11 associated with ATAC peaks within 20kb of the TSS, as well as the genes differentially regulated.

Subsequently, we sought to identify the degree of overlap between genes that are associated with ATAC peaks either in their introns or up to 20kb upstream of the TSS, and differentially expressed genes (DEGs) in each cell type (Fig. 6C-F). We found 152 DEGs with ATAC peaks in cranial osteoblasts, 251 in cartilage, 8 in *entpd5a*+ NSCs and 56 in intersegmental cells. In 3 out of 4 cell types, there were significantly more DEGs identified, compared to genes associated with ATAC peaks, an effect perhaps associated with the exclusion of peaks during differential accessibility analysis.

We then extracted the ATAC peaks associated with DEGs that have open chromatin in close proximity to the TSS. We examined the extracted peaks using the HOMER software, in order to identify enriched transcription factor binding sites in promoters of interest in each cell population. We intersected the resulting tables with RNA-seq datasets, identifying zebrafish orthologues of the predicted binding transcription factors that are differentially expressed in each given cell type (Table 2). Interestingly, we found that in osteoblasts there was both an enrichment for predicted binding sites of factors belonging to the Dlx family, and higher expression of orthologues of those particular members of the family compared to cartilage cells. Similarly, Hox family transcription factors were predicted to bind to open chromatin close to DEGs in *entpd5a*+ segments. Simultaneously, the zebrafish orthologues of the identified family members were differentially expressed in *entpd5a*+ segments. Finally, our analysis highlighted an enrichment of Fox and Sox family binding sites in both cartilage and intersegmental cells, accompanied by differential expression of the corresponding family member orthologues in our RNA-seq samples.

**Table 2.**
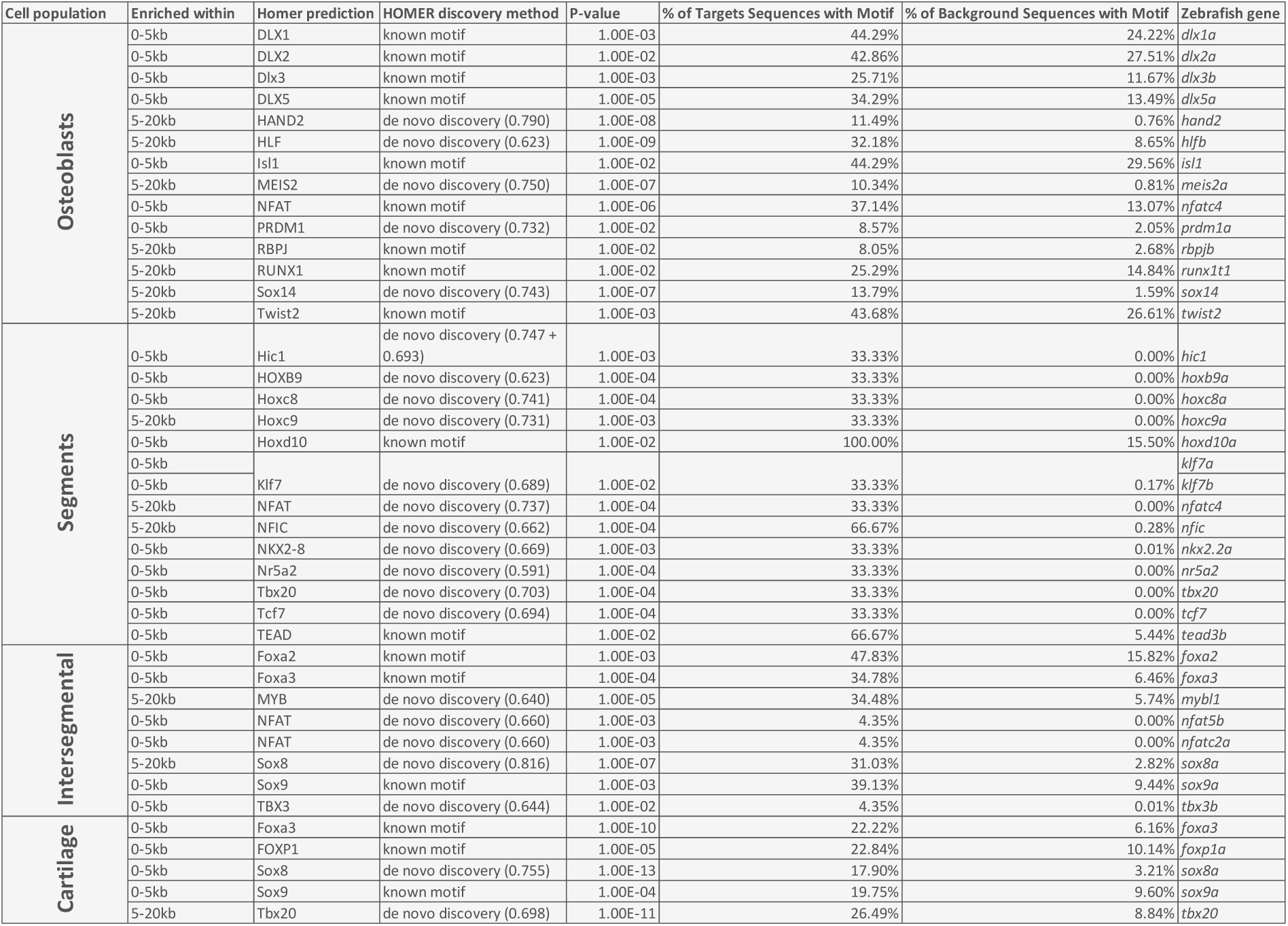
Candidate TFs regulating bone and cartilage development.

Finally, we used CIIIDER to search within introns 1 and 3, as well as within the 422bp element of the *entpd5a* promoter element identified in this work to play a crucial role in driving cranial osteoblast expression (Fig. 4A), for predicted binding sites of the classical and non-classical osteoblast regulators shown in Table 2 (Fig. 7). Notably, 12 putative Dlx factor binding sites were identified along the 422bp promoter region, with two of them contained entirely within the sequence deleted using the *entpd5a(r37):GFP* construct (Fig. 4A). The same area included a predicted Isl1 binding site in classical osteoblasts and two Hoxc8 binding sites in non-classical osteoblasts (Fig. 7). A total of six Runx2 binding sites were identified, of which two were positioned in the 422bp sequence. Of these, one site was found to drive expression in classical osteoblasts (Fig. 4). Although Runx2 was not one of the candidate factors identified in osteoblasts (Table 2), it is a factor known to play important regulatory roles in osteoblasts (Hojo et al., 2022; Lee et al., 2010; Nishio et al., 2006; Rashid et al., 2014). Moreover, our RNA-seq data showed that both *runx2a* and *runx2b* are slightly upregulated in osteoblasts versus cartilage (logFC = 1.39 and 2.64, respectively) and *runx2a* is upregulated in *entpd5a*+ segments versus intersegmental cells (logFC = 1.69). Compatible with this, our ATAC-seq data showed open chromatin in the vicinity of both genes, likely indicating open enhancers (data not shown).

**Figure 7.**
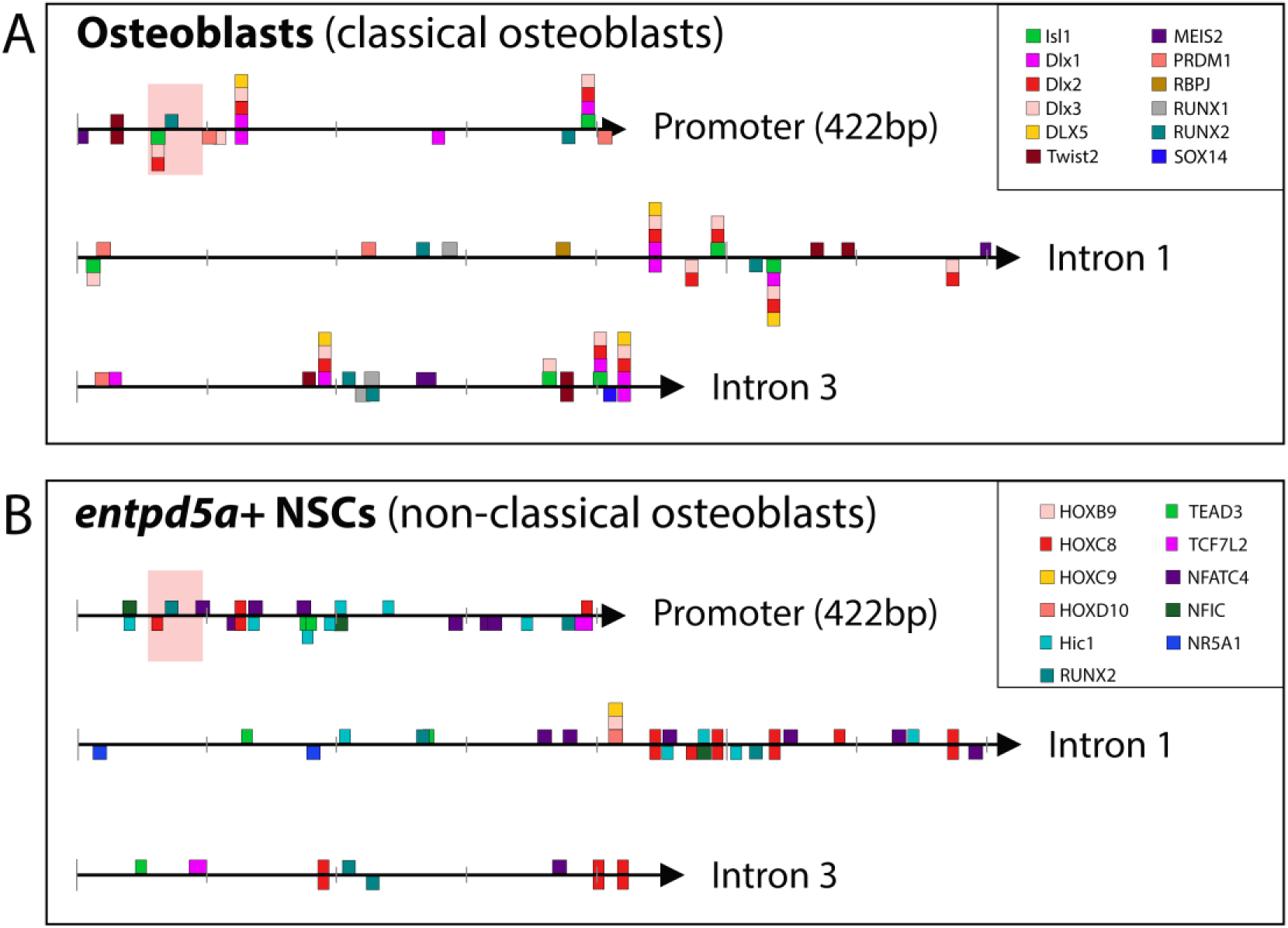
Distinct predicted binding sites could facilitate transcription factor binding of the *entpd5a* promoter, aiding cell type-specific regulation of the gene’s expression in classical and non-classical osteoblasts. Our analysis suggests that, with the exception of Runx2, distinct transcriptional regulators are functioning in (A) classical versus (B) non-classical osteoblasts. *In silico* analyses predict the presence and distribution of binding sites (squares, coloured according to transcription factor specificity) of the corresponding regulators in the 3 non-coding regions of *entpd5a* which are shown to have regulatory function. The width of each square represents the length of sequence of the respective binding site. The red box indicates the location of the 37bp deleted using *entpd5a(r37):GFP*.

In conclusion, we identified a number of strong candidates that are likely to function as transcriptional regulators in zebrafish classical and non-classical osteoblasts and that guide the dynamic and diverse expression of zebrafish *entpd5a* in these cell types.

## Discussion

In recent years, research on teleost fish identified the ability of the embryonic notochord to produce bone in a manner independent of classical sclerotome-derived osteoblasts, and to initiate the formation of mineralised chordacentra (Fleming et al., 2004; Lleras Forero et al., 2018). The cells responsible for this are non-classical osteoblasts within the notochord sheath, expressing the early bone marker *entpd5a* but not the classical osteoblast regulator *osterix* (Lleras Forero et al., 2018; Spoorendonk et al., 2008). Here, we aimed to better understand the genetic regulation and molecular signatures of classical and non-classical osteoblasts, and how those two populations differ from each other and from the closely related cranial chondrocytes and intersegmental NSCs. To this end, we characterised the newly generated transgenic line *entpd5a:Gal4FF*; *UAS:GFP* in combination with the previously described *R2col2a1a:mCherry* to isolate classical and non-classical osteoblasts as well as cartilage and intersegmental cells from the head and trunk of zebrafish larvae. We employed ATAC-seq to reveal the differentially accessible chromatin domains in *entpd5a*+ cells of head and trunk populations and performed RNA-seq in equivalent cell populations to understand the gene expression signatures accompanying observed chromatin changes. This first compilation of combined ATAC-seq and RNA-seq data from teleost osteoblasts enables the genome-wide analysis of osteoblast, chondrocytes, notochord sheath cells, and intervertebral disk precursors at both the chromatin and transcriptome level. This comprehensive dataset will be of use to a wide variety of researchers interested in osteogenesis, both within and outside the field of teleost developmental biology. For our immediate purposes, and as a proof of principle study, the data has allowed us to dissect the promoter organisation and to identify likely regulators of *entpd5a*, a critical gene in bone formation in zebrafish.

The ATAC-seq datasets of both *entpd5a*+ and *entpd5a*- cells in the head and trunk reveal the chromatin landscape in these skeletogenic cell populations. Although highly similar due to their respective origin from either cranial neural crest cells or notochord progenitor cells, differential chromatin accessibility analysis on head and trunk populations revealed putative enhancer regions of interest for bone vs. cartilage generation. Here, we validate our chromatin accessibility datasets and prove the significance of performing such analyses by demonstrating the regulatory importance of open regions detected both upstream of *entpd5a* and within the gene’s intronic regions, thus highlighting a highly complex promoter organisation. Specifically, we demonstrate that a series of active enhancers responsible for driving expression in classical osteoblasts are located in a short stretch of non-coding sequence within 2.2kb of the *entpd5a* ATG. Moreover, introns 1-3 contain regulatory regions required for *entpd5a* segmental activation in non-classical osteoblasts. Although we attempted to test several of the additional peaks found in non-coding areas surrounding the gene, more work is required to pinpoint the enhancers driving expression in cranial chondrocytes. In conclusion, the *entpd5a:Gal4FF* line has allowed us to refine the expression pattern of zebrafish *entpd5a* and to use ATAC-seq to elucidate the structure of its promoter.

We additionally performed RNA-seq in order to better understand the molecular signatures of the different cell populations and the functional significance of the observed chromatin configurations. As expected, our analyses reveal enrichment of bone-specific genes in *entpd5a*+ populations and, conversely, of cartilage-specific genes in *entpd5a*- populations. We went on to integrate the ATAC-seq and RNA-seq datasets to increase the predictive power of our analyses, which led to the determination of candidate transcriptional regulators for each cell type of interest. Our results suggest functional roles of Dlx family genes in classical osteoblasts, of Hox family regulators in non-classical osteoblasts, and of Foxa and Sox family members in *entpd5a*- cells of both the head and the trunk. It thus appears that gene expression is not regulated in the same manner in classical and non-classical osteoblasts, since the identified candidate regulators showed no overlap. In contrast, chondrocytes of the head and intersegmental NSCs demonstrated a surprising amount of overlap in putative regulators governing gene expression specific to their skeletogenic roles.

Identification of putative functions of Dlx family transcription factors in classical osteoblasts is compatible with the long-recognised roles of these factors in upregulating key osteoblast genes during all stages of osteoblast specification and maturation (Hassan et al., 2004; Li et al., 2008; Shirakabe et al., 2001). Significantly, Dlx factors have been implicated in recruiting Osterix, the master regulator of osteoblasts, to osteoblast enhancers in mice (Hojo et al., 2016). Interestingly, it has been shown that distinct Dlx family members become active in different stages of osteoblast development, with chick Dlx3 being able to activate mature osteoblast expression programs (Li et al., 2008). Consistent with this, our analyses reveal enrichment of *dlx3b* expression in zebrafish classical osteoblasts. In non-classical osteoblasts, we identified several Hox factors amongst predicted regulators of skeletal fate genes. Hox family genes have long been known for their crucial roles in skeletal patterning (Fromental-Ramain et al., 1996a, 1996b; Wellik and Capecchi, 2003), and in the adult skeleton (Song et al., 2020). In future work, the binding affinity of zebrafish Dlx and Hox factors on the promoters of osteoblast genes such as *entpd5a*, as well as generating mutants in specific *dlx* and *hox* genes would be appropriate means to obtain further insight into the gene regulatory network governing osteogenesis and axial patterning.

In regards to candidate regulators with putative roles in *entpd5a*- cells, Foxa2 and Foxa3 are transcription factors with confirmed roles in chondrogenesis (Ionescu et al., 2012). Furthermore, SOX9 is known as the master regulator of chondrocyte differentiation, and competition between SOX9 and FOXA have been shown to aid this process (Tan et al., 2018). Both *entpd5a*- cell populations investigated here differentially express the zebrafish genes *foxa2*, *foxa3* and *sox9a*, while the second SOX9 orthologue, *sox9b*, is enriched in cranial *entpd5a*- cells. We thus postulate that the mechanism of chondrocyte differentiation is likely conserved between amniotes and zebrafish.

In summary, integration of ATAC-seq and RNA-seq has been valuable in predicting candidate regulators for skeletogenic cells found in teleost fish in a non-biased manner. Although this should not be viewed as a comprehensive list of regulators, it provides guidance in the first steps of elucidating genetic interactions taking place in these cell types, and particularly in the non-classical osteoblasts of the notochord, which have only been recently started to be analysed.

In addition to its use in better understanding the gene regulatory programs acting in skeletogenic cells, our approach of performing both RNA-seq and ATAC-seq using *in vivo*-derived samples and analysing the data not only individually but also in an integrated manner, provides a comprehensive resource that has hitherto not been available. Although osteoblasts of mammals and birds have been the subject of transcriptomic analyses for decades, the vast majority of RNA-seq and ATAC-seq studies have been performed using *in vitro*-differentiated cells (Hao et al., 2022; Liu et al., 2022; Yu et al., 2021). To our knowledge, only one study thus far used *in vivo*-derived mouse neonatal skeletal cells to perform both assays (Hojo et al., 2022). Furthermore, RNA-seq and ATAC-seq studies on osteoblasts of teleost fish remain extremely limited. With the exception of three studies: one in which scRNA-seq was performed on the zebrafish adult regenerating fin (Tang et al., 2022), another study where RNA-seq was performed on medaka osteoblasts (Phan et al., 2020), and a recent work on the transcriptomic signatures of classical osteoblasts isolated from zebrafish embryos at 4 dpf (Raman et al., 2024), we are not aware of RNA-seq and ATAC-seq analyses that have been performed on embryonic classical osteoblasts of teleost fish. Thus, our study fills a void and moves teleost osteoblasts on par with the single mouse study (Hojo et al., 2022) that provided both ATAC-seq and RNA-seq data.

Of note, the significance of studying the molecular signatures of osteoblasts in teleost fish has become more pressing in recent years. It has been observed that ‘aquatic’ and ‘amphibian’ osteoblasts and cartilage behave differently to ‘terrestrial’ osteoblasts in terms of gene expression (Nguyen and Eames, 2020). Specifically, the osteoblasts of teleost fish and amphibians, whose characteristics are putatively closer to a more ancestral state of skeletal differentiation compared to amniotes, appear to share gene expression with chondrocytes, a pattern which we also observed in our datasets, while amniote osteoblasts have suppressed chondrocyte markers in the course of evolution. Despite this, the vast majority of NGS studies have been carried out on osteoblasts of amniotes, leading to the establishment of an incomplete picture of the osteoblast (Nguyen and Eames, 2020). In this work, we show that the bone marker *entpd5a* is expressed in chondrocytes of the developing jaw, in a manner similar to *osterix* (Hammond and Schulte-Merker, 2009). Furthermore, we show that several other genes conventionally thought of as cartilage-specific, such as *ccn6, sox9a, acana* and *col9a2* are expressed at low levels in osteoblasts. These discoveries reinforce the arguments regarding the chondrocyte origin of osteoblasts, while also pointing towards cells that might be prone to transdifferentiate into osteoblasts (Hammond and Schulte-Merker, 2009; Thesingh et al., 1991; Zhou et al., 2014).

Moving beyond the classical osteoblasts, we also here provide means to investigate non-classical osteoblasts, a cell type confined to teleosts that has recently been shown to play crucial roles in the initiation of spine development (Grotmol et al., 2003; Lleras Forero et al., 2018; Renn et al., 2013). Very little is known to date about the mechanisms of gene regulation in these non-classical osteoblasts, limited to RNA-seq data that indicate a role for BMP and Notch signalling in patterning the cells of the notochord sheath (Wopat et al., 2018; Peskin et al., 2023). However, the gene regulatory interactions that are required to bring about the segmental expression of *entpd5a* remains unknown. We here demonstrate and experimentally validate a short piece of promoter sequence to be sufficient in governing *entpd5a* expression in NSCs, setting the stage for identifying the gene regulatory network that determines transcriptional control of this key factor in axial skeletogenesis.

Finally, this work provides insight into the mechanism of regulation of the early bone marker *entpd5a*, which we show is regulated in an unexpectedly dynamic manner throughout zebrafish embryonic development. *entpd5a* is not only of great interest because it acts as a specific osteoblast marker, but also because it is non-dispensable for bone development in zebrafish (Huitema et al., 2012). Importantly, *entpd5a* expression presents in a segmented manner along the zebrafish embryonic notochord, predetermining the locations of chordacentra mineralisation and future vertebrae position. This segmentation was shown to be independent of the somitic segmentation clock (Lleras Forero et al., 2018), arguing for an independent manner of generating the periodic appearance of *entpd5a* expression within NSCs. The molecular mechanism guiding this stereotypic pattern remains unclear (Fernández Arancibia et al., 2022).

Here, we show that *entpd5a* introns 1-3, together with enhancers located upstream of the ATG, drive segmental notochord expression of *entpd5a*. Furthermore, we used our NGS analyses to predict candidate regulators, including *dlx*, *hox* and *runx* family genes, putatively binding on the identified enhancer sites to drive *entpd5a* expression in both classical and non-classical osteoblasts. Importantly, we identified a 37bp region within 2.2kb of the ATG, predicted to be able to bind Runx2a or Runx2b, as well as various members of the *dlx* gene family. *runx2* family members are of particular interest to the process of osteogenesis, as mammalian Runx2 is known to play important roles in the regulation of osteoblast genes, both through its action as a pioneer factor, playing a crucial role in the plasticity between chondrocyte and osteoblast fates (Hojo et al., 2022), and through its interaction with co- factors to directly upregulate osteogenic genes (Lee et al., 2010; Nishio et al., 2006; Rashid et al., 2014). Although *runx2a* and *runx2b* were not thought to be expressed in mature osteoblasts in teleost fish (Huycke et al., 2012), via RNA-seq data we detect low level expression of at least one of the two factors in *entpd5a*+ cells of the head and of the trunk, rendering them plausible *entpd5a* regulators in classical and non-classical osteoblasts. Our *in silico* predictions and presented hypotheses are to be validated by functional analyses (such as generating mutants), and this information can be built on to reveal gene regulatory network upstream of *entpd5a* in both types of zebrafish osteoblasts. Moreover, the role of Entpd5 in bone formation should be determined in other species and the evolutionary conservation of the mechanisms which we begin to identify here should be confirmed.

Overall, we here characterise the organisation of the complex promoter of *entpd5a*, and identify critical genomic elements that govern transcriptional control of this early and highly specific marker of classical and non-classical osteoblasts. En route, we suggest candidate transcriptional regulators that can now be tested. We furthermore provide a resource with a wide range of possible uses in the field, making both ATAC-seq and RNA-seq data sets available that are based on chondrocytes and osteoblasts isolated from *in vivo* derived material.

## Materials and Methods

### Fish raising and husbandry

Zebrafish (*Danio rerio*) strains were maintained according to FELASA recommendations (Aleström et al., 2020). Animal experiments have been performed according to animal ethics committees’ guidelines at the University of Münster, Germany. Tissues were collected according to the approved protocol for tissue removal. Embryonic developmental stages were determined according to Kimmel et al., 1995 (Kimmel et al., 1995). In this study, the following published transgenic lines have been employed: *TgBAC(entpd5a:pkRed)^hu7478^*and *TgBAC(entpd5a:Kaede)^hu6867^* (Lleras Forero et al., 2018), *Tg(UAS:GFP)^nkuasgfp1a^* (Asakawa et al., 2008), *Tg(R2col2a1a:mCherry)^ue401Tg^*(Lopez-Baez et al., 2018), *TgBAC(entpd5a:Gal4FF)^mu412^* (Labbaf et al., 2022).

### Generation of constructs and transgenic lines

For generation of all lines, we amplified by PCR promoter sequences from zebrafish genomic DNA. The transgenic line *Tg(-2.2entpd5a:GFP)^mu416^* (referred to as *entpd5a(2.2):GFP*) was generated by insertion of the PCR-amplified promoter sequence into a miniTol2-GFP vector via an EcoRI restriction site located directly upstream of the GFP. The line *Tg(-2.2entpd5a:GFP-introns)^mu419^*(referred to as *entpd5a(2.2-introns):GFP*) and the injected construct -2.2*entpd5a*:*GFP-coq6* (referred to as *entpd5a(2.2-coq6):GFP*) were both generated by inserting PCR-amplified fragments downstream of the GFP cassette of the *entpd5a(2.2):GFP* vector via an EcoRV restriction site. *Tg(-5.7entpd5a:GFP)^mu418^*(referred to as *entpd5a(5.7):GFP*) was generated in a 2-step process, first using NEBuilder® HiFi DNA Assembly (New England Biolabs, E2621S) with the *entpd5a(2.2):GFP* construct, digested with MscI restriction enzyme, and 2 PCR-amplified fragments of 1023bp and 1040bp, respectively, as templates. The additional fragments were thus consecutively inserted directly upstream of the 2.2kb sequence. Subsequently, an 1.4kb PCR-generated fragment was inserted via a BamHI restriction site directly upstream of the previously inserted *entpd5a* promoter elements. All primers used for cloning are listed in Supplementary Table 2.

The deletion constructs shown in Fig. 4 were generated by PCR amplification of the *entpd5a(2.2):GFP* vector using phosphorylated primers listed in Supplementary Table 2. The PCR constructs were self-ligated to generate injection vectors. All lines were generated by injecting 25pg of plasmid DNA and 25pg of Tol2 transposase mRNA into single cell-stage embryos.

The *TgBAC(Δ21entpd5a:pkRed)^mu420^* and *TgBAC(Δ10entpd5a:pkRed)^mu420^*lines (referred to as *entpd5a(Δ21):pkRed* and *entpd5a(Δ10):pkRed,* respectively) were generated using BAC recombineering based on the protocol previously described (Bussmann and Schulte-Merker, 2011). Briefly, *Escherichia coli* carrying the *entpd5a:pkRed_Tol2* BAC construct were electroporated with a PCR-amplified sequence in which the open reading frame (ORF) of the antibiotic Spectinomycin is positioned downstream of the Kanamycin promoter. Flanking the Kan/Spectinomycin sequence were homology arms matching either side of the stretch of non-coding sequence targeted for deletion. The Spectinomycin ORF was thus inserted in the *entpd5a:pkRed* construct, replacing 21kb of non-coding sequence upstream of the ATG of *entpd5a* (*entpd5a(Δ21):pkRed*), or 10kb of coding and intronic sequences (*entpd5a(Δ10):pkRed*). 100pg of modified BAC DNA were injected into single cell-stage embryos, together with 25pg Tol2 transposase mRNA.

### Imaging and image analysis

Confocal imaging was performed using a Leica SP8 microscope (10x and 20x objectives) employing Leica LAS X 3.5.6.21594 software (https://www.leica-microsystems.com/). z-stacks for the 20x objective were taken at an interval of 2.5μm. Stacks were flattened by maximum projection, images were processed and fluorescent measurements were performed using Fiji (Fiji Is Just ImageJ) (Schindelin et al., 2012). Adobe Photoshop CC 2022 was also used for image processing. Photo- conversion was performed on embedded *TgBAC(entpd5a:Kaede)* embryos via exposure for 5min to UV light. Quantification of fluorescence (Corrected Total Cell Fluorescence; Figure 2) was performed using the following equation according to instructions in https://theolb.readthedocs.io/en/latest/imaging/measuring-cell-fluorescence-using-imagej.html:

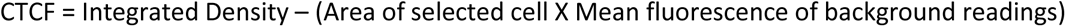

### FACS and RNA-sequencing

Embryos at 15 dpf were collected, anaesthetised and heads were removed through a clean cut directly anterior to the swim bladder. The procedure does not allow for a complete separation of notochordal non-classical osteoblasts from cranial classical osteoblasts, as the notochord extends into the cranium. However, the amount of sheath cells in that portion of the notochord is negligible, compared both to the number of classical (cranial) osteoblasts in head samples, and to notochord cells isolated in trunk samples. Heads and trucks were collected separately and incubated for 1hour at 34°C, at 400rpm in 0.25% Trypsin/EDTA in PBS (VWR, L0910-100) containing Collagenase B (Roche, 11088831011; 1% for head dissociation and 2% for trunk dissociation). Upon full dissociation, cells were centrifuged at 1,400rpm for 5min at room temperature. The supernatant was discarded and cells were washed with 1ml L15 medium (Sigma-Aldrich, L5520)/10% FCS solution. The wash step was repeated and samples of the same tissue were combined into a single Eppendorf tube in 1ml L15/FCS wash solution. Upon centrifugation and removal of the supernatant, pellets were resuspended in 1ml wash solution. Cells were passed through a 50μm and subsequently a 30μm strainer, and transferred to a FACSAria IIIu cell sorter (BD Biosciences) on ice. For FACS sorting the FACSDiva 8.0.1 (https://www.bdbiosciences.com/) and FlowJo 10.6.1 (https://www.flowjo.com/) software was used. For ATAC-seq and RNA-seq samples the 100μm and the 70μm nozzle were used, respectively. For ATAC-seq cells were sorted into ice-cold L15/FCS medium at 4°C, while for RNA-seq cells were collected directly into ice-cold RSB/β-Mercaptoethanol lysis buffer (QIAGEN RNeasy Plus Micro Kit, 74034; prepared according to manufacturer’s instructions).

For RNA-sequencing, RNA was isolated using the RNeasy Plus Micro Kit (QIAGEN, 74034). Samples were sent to Eurofins Genomics for library preparation and paired-end sequencing with the INVIEW Transcriptome Ultra Low service (2x 150). Per sample, 15M reads were sequenced. De-multiplexed sample files were obtained.

### ATAC-sequencing

ATAC-seq was performed according to previously published protocols (Buenrostro et al., 2015), with the following modifications. Cells were incubated in lysis buffer (10mM Tris-HCl, pH 7.4, 10mM NaCl, 3mM MgCl2, 0.1% Tween20, 0.1% NP40, 0.01% Digitonin) for 3min on ice. Lysis was stopped by resuspending in ice-cold wash buffer (10mM Tris-HCl, pH 7.4, 10mM NaCl, 3mM MgCl2, 0.1% Tween20) prior to centrifugation. Depending on the numbers of cells isolated in each sample, we adjusted the volume of enzyme added to our transposition reaction (1x Tagment DNA Buffer, 0.01% Digitonin, 0.1% Tween20, TDE1 enzyme) and the length of transposition on a heat block set at 37°C shaking at 1,000rpm (Table 3). The Tagment DNA Buffer and TDE1 enzyme are both contained in the Tagment DNA TDE1 Enzyme and Buffer Kit (Illumina, 20034197). The transposed chromatin was isolated using the Minelute PCR Cleanup kit (Qiagen, 28004).

**Table 3.** Volume of TDE1 Tagment DNA Enzyme, total reaction volume and incubation time of transposition reaction according to cell numbers.

| Cell number | 5-6.9k | 7-9.9k | 10-12.9k | 13-20.9k | 21-39.9k | 40-60k |
| --- | --- | --- | --- | --- | --- | --- |
| TDE1 enzyme | 1µl | 1.5µl | 1.8µl | 2.3µl | 3µl | 3µl |
| Total reaction volume | 20µl | 20µl | 50µl | 50µl | 50µl | 50µl |
| Incubation time | 30min | 30min | 30min | 30min | 30min | 45min |

All PCR amplification and quality control steps were carried out according to Buenrostro et al., 2015 (Buenrostro et al., 2015). Following amplified DNA purification, AMPure XP SPRI beads (Beckman Coulter, A63881) were used (1.4x sample volume) to remove primer dimers from the chromatin sample. The beads were brought to room temperature and thoroughly vortexed before use. The sample plus beads mixture was incubated at room temperature for 5min, then placed on the magnetic stand for 5min to clear the solution. The supernatant was discarded and the beads were washed for 30sec with 80% ethanol. The wash step was repeated. Following removal of the supernatant, the beads were air-dried, removed from the magnetic stand and 20μl of EB buffer was added for DNA elution. The sample was mixed and incubated for 2min at room temperature, then placed back on the magnetic stand for 5min. The supernatant was transferred to a fresh tube.

The quality of purified chromatin was assessed using the Bioanalyzer High Sensitivity DNA Analysis kit (Agilent). Libraries were sequenced using the NextSeq 500/550 Mid Output Kit v2.5 (150 Cycles) (Illumina, 20024904) on the sequencing platform Illumina NextSeq500. 20M reads were sequenced per sample.

### Sequencing Data analysis

For ATAC-seq, sequences were de-multiplexed into fastq files applying standard parameters of the Illumina pipeline (bcl2fastq) using Nextera index adapters. The individual fastq files were processed with fastp (version 0.20.0) (Chen et al., 2018) and mapped to the zebrafish reference genome GRCz11 using Bowtie2 (version 2.3.5.1) (Langmead and Salzberg, 2012) with the ‘very-sensitive option’. Sequences with mapping quality >30 were isolated and sorted using SAMtools (version 1.9) (Danecek et al., 2021). Mitochondrial reads and duplicates were removed using BEDTools (version 2.28.0) (Quinlan and Hall, 2010) and Sambamba (version 0.7.1) (Tarasov et al., 2015), respectively. Reads were shifted using the standard method for ATAC samples and bigwig files were generated for visualisation using deepTools (version 3.3.1) (Ramírez et al., 2016). IGV (https://igv.org/app/) was employed for peak visualisation. MACS2 (version 2.2.6) (Zhang et al., 2008) was used to call peak enrichment with the settings -g 1.412e9 --keep-dup all --nomodel --shift -100 --extsize 200. ataqv (version 1.3.0) (Orchard et al., 2020) was employed for assessing the fragment distribution analysis and TSS enrichment scores. To determine differentially accessible chromatin regions and produce plots, we used the R package DiffBind (version 3.8.4) (Ross-Innes et al., 2012). The summits parameter was set to 150, to account for peak read lengths of 301bp. Head samples (osteoblasts vs. cartilage) and trunk samples (*entpd5a*+ segments vs. intersegmental regions) were individually contrasted. An element was considered *entpd5a*-positive if log2 fold change in read density was ≤-1 and FDR≤0.05. Conversely, *entpd5a*-negative elements were log2 fold change ≥1 and FDR≤0.05.

RNA-seq analysis was carried out by first pre-processing reads and removing duplicates using fastp (version 0.23.2) (Chen et al., 2018), aligning the cleaned reads to the zebrafish reference genome GRCz11 using the STAR aligner (version 2.7.2b) (Dobin et al., 2013), isolating and sorting high mapping quality reads (q > 30) using SAMtools (version 1.9) (Danecek et al., 2021). Raw read counts for each gene were calculated using the function featureCounts from the Subread package (Liao et al., 2014). The R package DESeq2 (version 1.38.3) (Love et al., 2014) was employed for differential expression analysis with an adjusted p-value cut-off of ≤0.05. As for ATAC-seq, *entpd5a*+ and *entpd5a*- populations from head and trunk samples were contrasted amongst each other, and only genes with |log2FC|>0.5 were considered for further analyses. PCA plots were produced with DESeq2, and heatmaps using the package ComplexHeatmap (Gu, 2022; Gu et al., 2016).

### Integration of ATAC-seq and RNA-seq and binding site prediction

Data integration was performed using the R package tidyverse (Wickham et al., 2019). Visualisations were generated using the R packages VennDiagram (Chen and Boutros, 2011) and ggplot2. HOMER software was used to identify enrichment in transcription factor binding sites (Heinz et al., 2010). The program CIIIDER was used to identify binding sites within the *entpd5a* promoter sequences (Gearing et al., 2019).

### Statistics, graphs and reproducibility

All images shown in the figures are representative examples of the respective expression patterns. All statistics were performed using R. For visualisations, we used the R package ggplot2.

## Data availability

Processed as well as unprocessed ATAC-seq and RNA-seq data generated as part of this work can be accessed via Gene Expression Omnibus (accession numbers GSE267332 and GSE267333, respectively). Furthermore, code used in data processing has been deposited, and is freely available on Github (https://github.com/KleioP/ATAC_RNAseq_analysis/tree/Petratou-et-al.%2C-2024).

## Acknowledgments

We thank Dr. Backialakshmi Dharmalingam, Dr. Kishor K. Sivaraj and Katja Müller (MPI Münster) for support with performing ATAC-seq, as well as Paul-Georg Majev (MPI Münster) and Dr. Ada Jimenez-Gonzalez (University of Birmingham) for their valuable feedback on analysing NGS data.

---

**Supplementary Figure 1.**
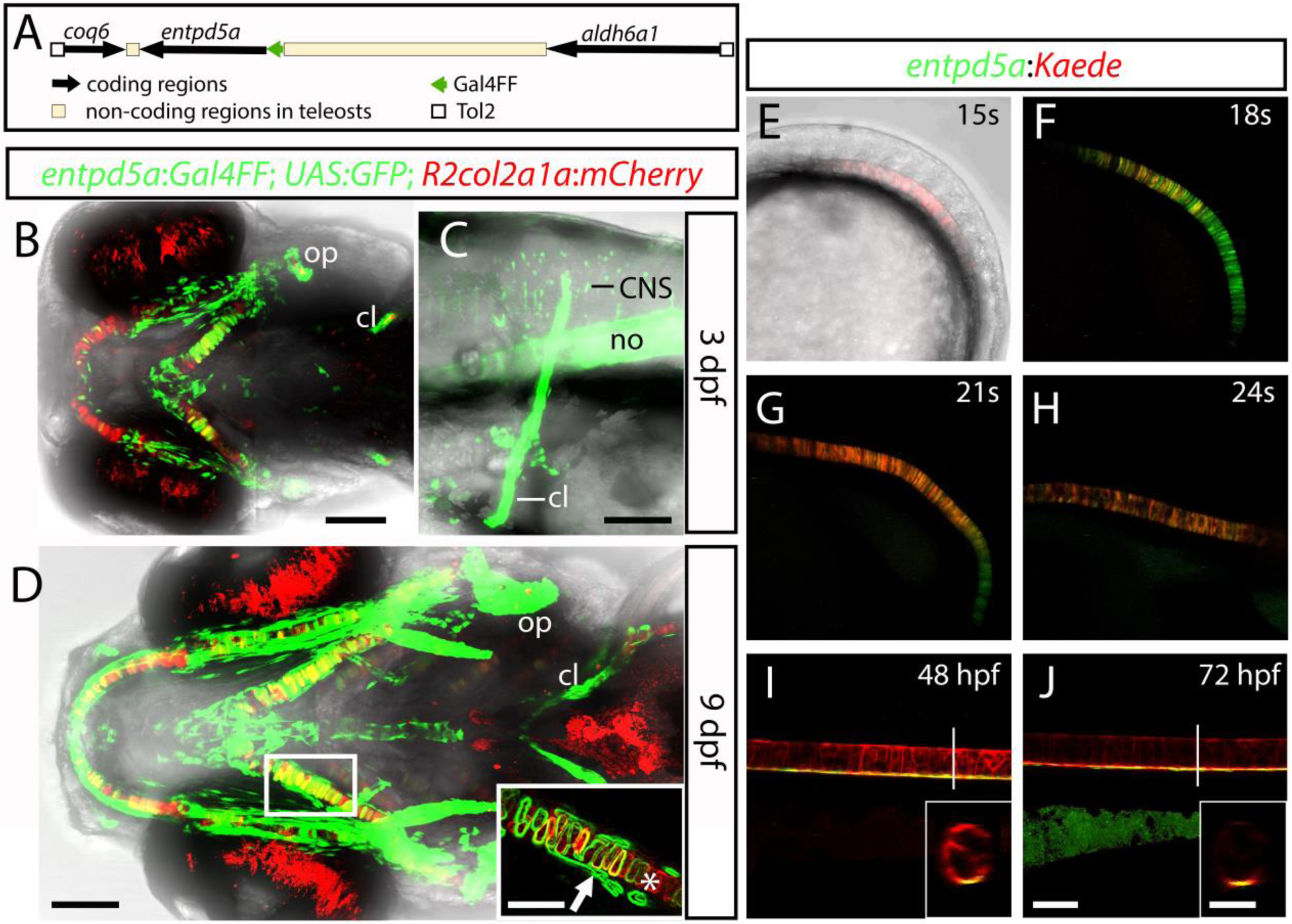
The complex expression pattern of *entpd5a* is dynamically regulated during zebrafish development. (A) Schematic of the BAC driving expression of Gal4FF in the *entpd5a:Gal4FF* transgenic line. The BAC contains the entire open reading frame of *entpd5a*, and part of the open reading frames of adjacent genes, *coq6* and *aldh6a1*. (B, C, D) GFP expression is detected in osteoblasts (arrow) and (partially) in cartilage (asterisk; D, inset) making up the head skeleton. (C) Strong GFP expression is seen in the notochord and the cleithrum, but also in a subset of CNS neurons. (E-J) Using the *entpd5a:Kaede* photoconversion line we first detect *entpd5a* expression at the 15 somite-stage (E). Following the same embryo, active expression of the gene continues until prior to 24 hpf (F-H). Between 24 hpf and until notochord segmentation takes place, *entpd5a* is only actively expressed in the ventral-most NSCs of the notochord, while expression in the remaining NSCs and vacuolated cells is turned off (I, J). (I-J, insets) Cross section of the notochord at 48 hpf and at 72 hpf in the position indicated by the white vertical line. (B) ventral view; (C-J) lateral views, anterior towards the left. (B-J) scale bars: 100μm. (D, I, J) insets scale bars: 50μm.

**Supplementary Figure 2.**
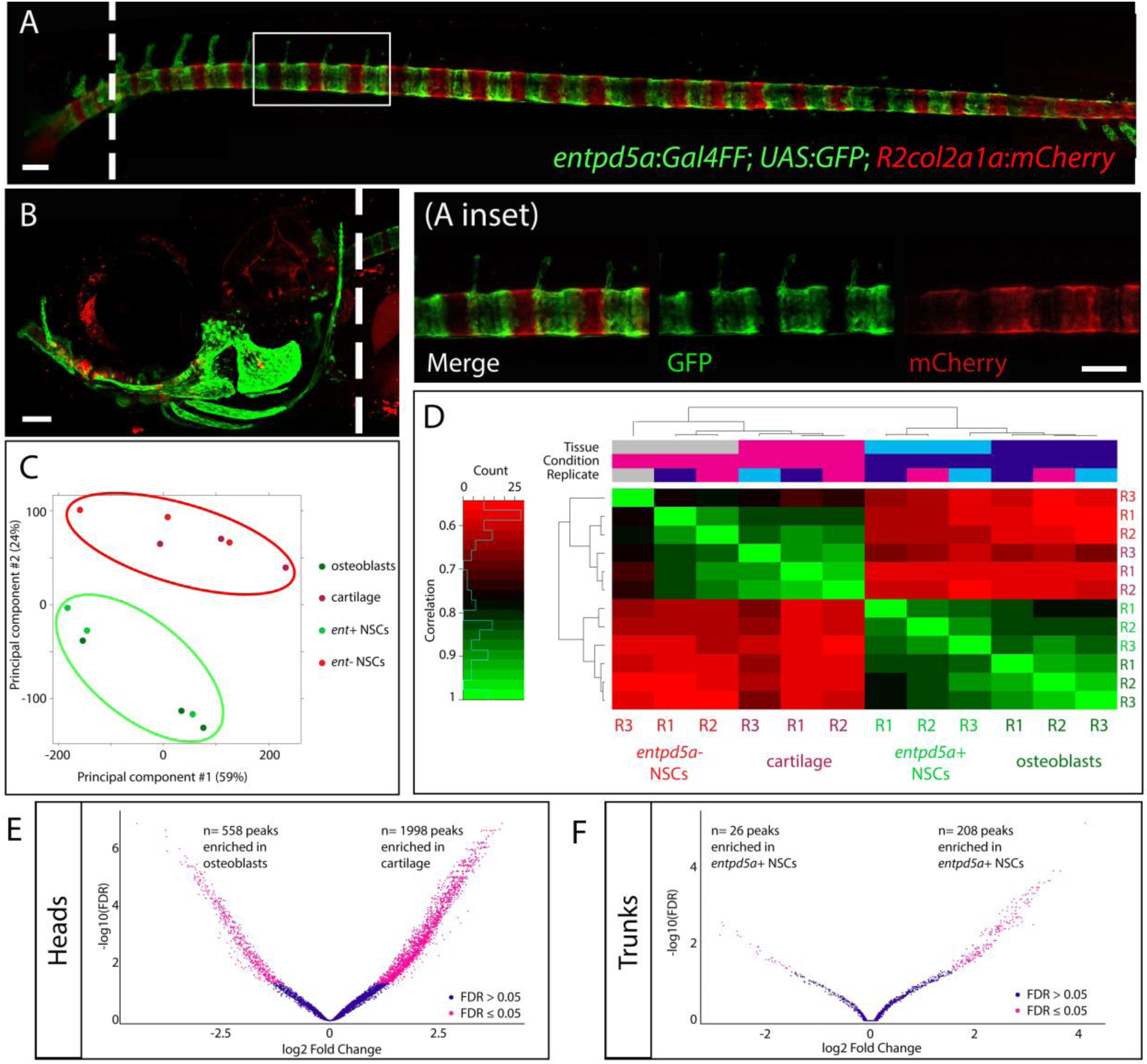
Quality assessment of ATAC-seq of skeletogenic cells. (A, B) Trunk and head, respectively, of transgenic fish at 15 dpf, used for collection of FAC-sorted cells. Dashed lines indicate the site where the cut was made to separate head from trunk tissue. GFP+ cells indicate mineralising cells (A, NSCs; B, cranial osteoblasts) while mCherry+ cells indicate (A) intersegmental NSCs and (B) head cartilage. (C) PCA analysis, with red circle indicating mCherry+ cell samples and green circle GFP+ cell samples. (D) Heatmap indicating correlation amongst individual replicates. (E, F) Volcano plots showing for (E) head and (F) trunk the identified significantly accessible chromatin regions. Positive log Fold Change indicates regions open in mCherry+ cells, negative log Fold Change indicates regions open in GFP+ cells. (A, B) lateral views, anterior to the left. Scale bars: 100μm.

**Supplementary Figure 3.**
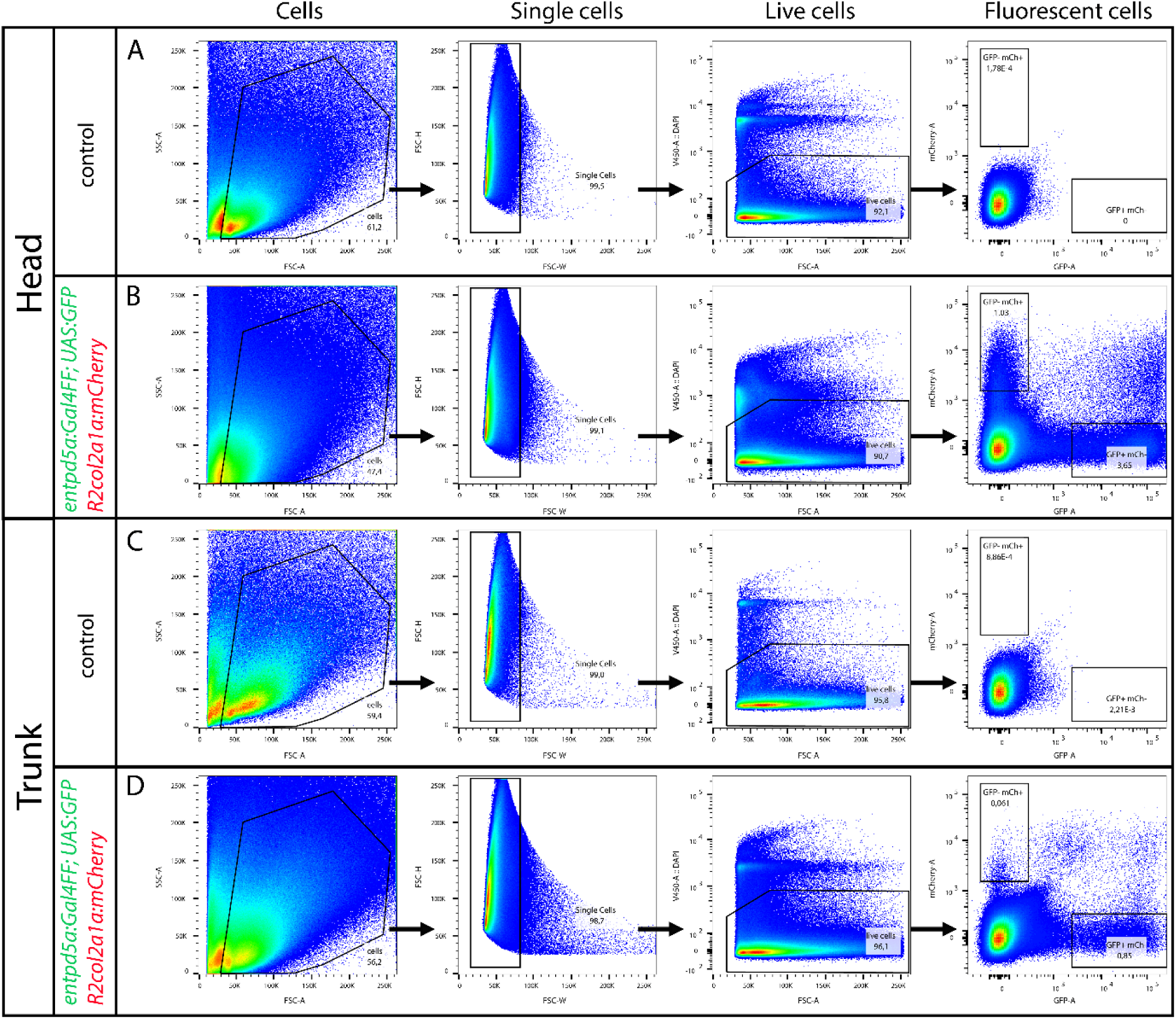
Gating strategy for FAC-Sorting cells for ATAC-seq and RNA-seq. Cells from (A) heads and (C) trunks of GFP-; mCherry-embryos were used as gating controls. Cells from (B) heads and (D) trunks of GFP+ ; mCherry+ siblings were then sorted.

**Supplementary Figure 4.**
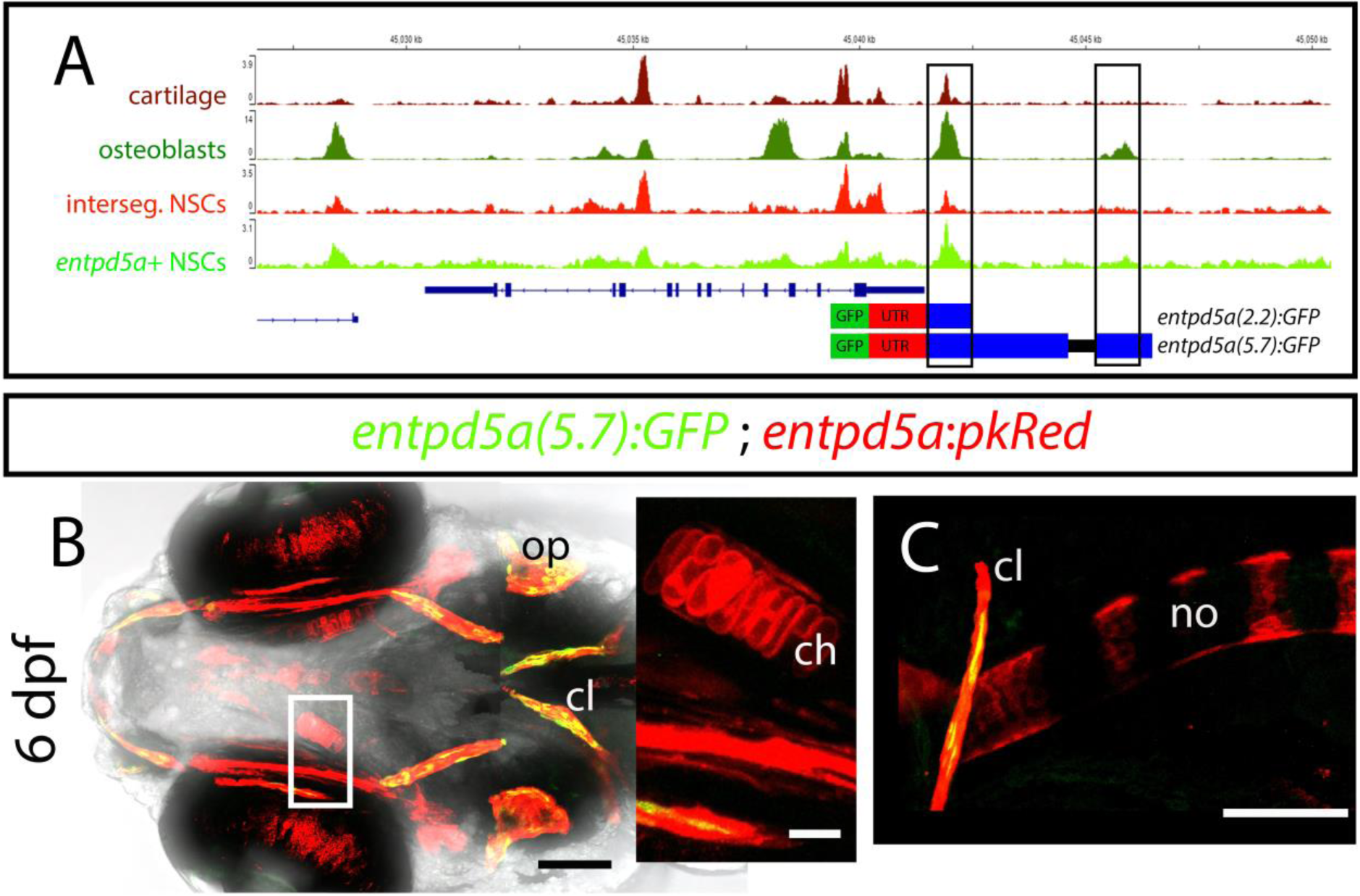
No additional enhancers are present in the sequence within peak number 5. (A) Chromatin accessibility profiles of representative samples of each cell type. In squares are the peak region covered by the *entpd5a(2.2):GFP* construct and the one covered by the *entpd5a(5.7):GFP* construct. (B, C) GFP driven in a stable line by the 5.7kb upstream of the start codon of *entpd5a*, is only expressed in cranial osteoblasts (B), but not in the notochord (C) or in cartilage of the head skeleton (B, inset). (B) ventral view; (C) lateral view. Anterior towards the left. (D, E) scale bars: 100μm. (D inset) scale bar: 20μm.

**Supplementary Figure 5.**
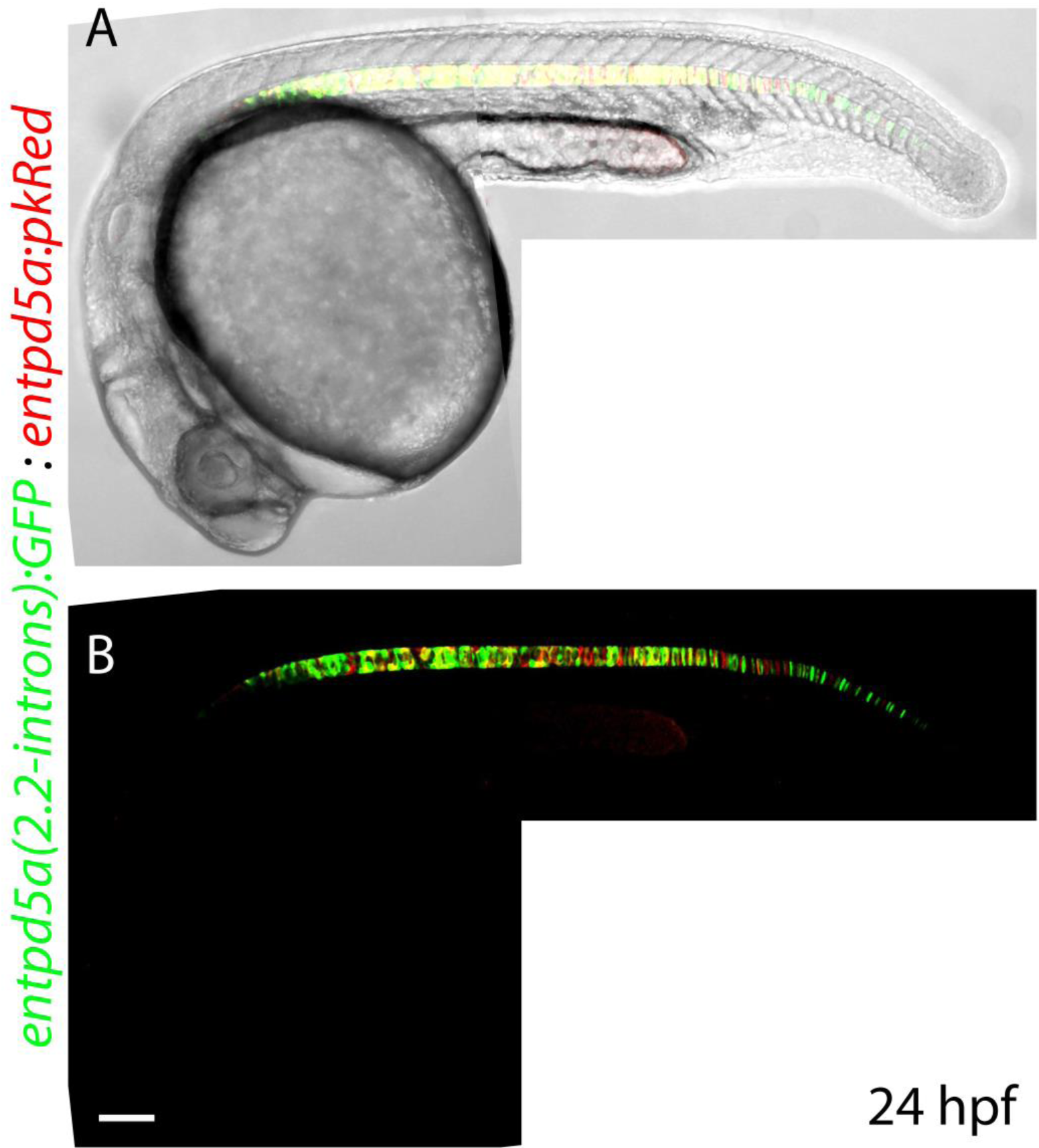
GFP expression in the notochord under the control of the introns is first detected in notochord progenitor cells. (A, B) GFP and pkRed expression (under the control of the complete BAC) appear to completely overlap in notochord progenitor cells at 24 hpf. Scale bar: 100μm.

**Supplementary Figure 6.**
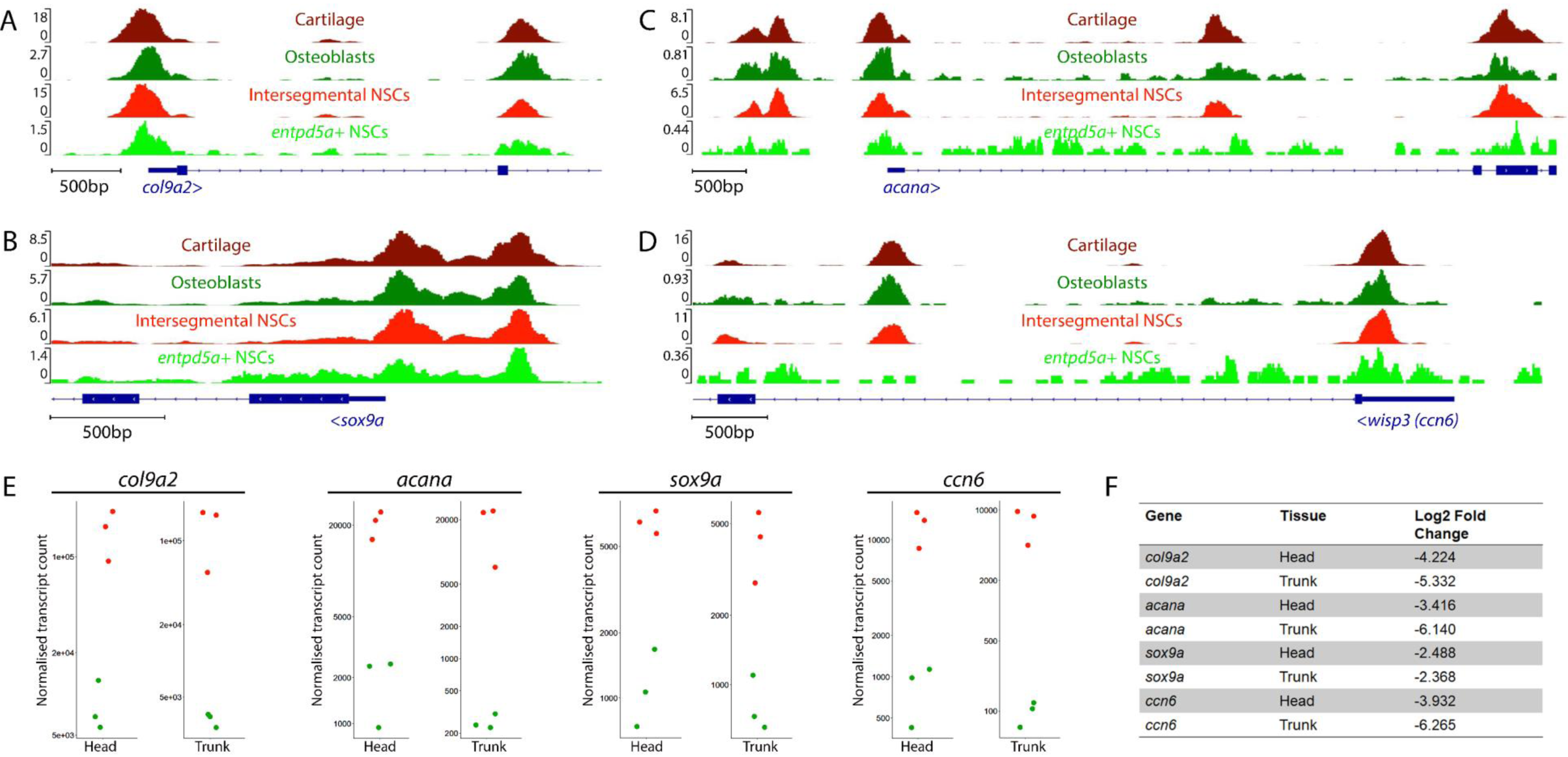
Chondrocyte markers maintain an open chromatin configuration in osteoblasts and are actively expressed in *entpd5a*+ cells in levels lower than that found in *entpd5a*-cells. (A-D) Views of the *col9a2* (A), *sox9a* (B), *acana* (C) and *ccn6* (D) promoters in representative ATAC samples for each of our cell populations, with the scale bars indicating the maximum normalised ATAC signal (read counts) in each sample. (E) Normalised *col9a2*, *sox9a*, *acana* and *ccn6* transcript counts in the three *entpd5a*+ (green spots) and the three *entpd5a*-(red circles) samples sequenced in the head and in the trunk. (F) Corresponding log2 Fold Change values for each gene when *entpd5a*+ vs. *entpd5a*-cell populations are compared within each tissue.

## Supplementary tables

**Supplementary Table 1.** Quality control measurements for ATAC-seq samples. Fragment Length Distribution Distance (FLDD) is defined as the distance of the experiment’s FLD to a reference distribution. Negative and positive distances are associated with under- and over-transposed samples, respectively. TSS enrichment score is calculated based on the normalised fragment coverage +/- 2kb around the TSS.

| Sample | FLDD | TSS score | Sample | FLDD | TSS score |
| --- | --- | --- | --- | --- | --- |
| Osteoblasts rep1 | 0,267 | 2,332 | *entpd5a*+ NSCs rep1 | 0,259 | 2,168 |
| Osteoblasts rep2 | 0,207 | 4,996 | *entpd5a*+ NSCs rep2 | 0,258 | 2,479 |
| Osteoblasts rep3 | 0,219 | 4,236 | *entpd5a*+ NSCs rep3 | 0,231 | 4,245 |
| Chondrocytes rep1 | 0,211 | 6,177 | *entpd5a*- NSCs rep1 | 0,299 | 3,511 |
| Chondrocytes rep2 | 0,215 | 4,244 | *entpd5a*- NSCs rep2 | 0,182 | 4,507 |
| Chondrocytes rep3 | 0,230 | 3,020 | *entpd5a*- NSCs rep3 | 0,181 | 2,097 |

**Supplementary table 2.**
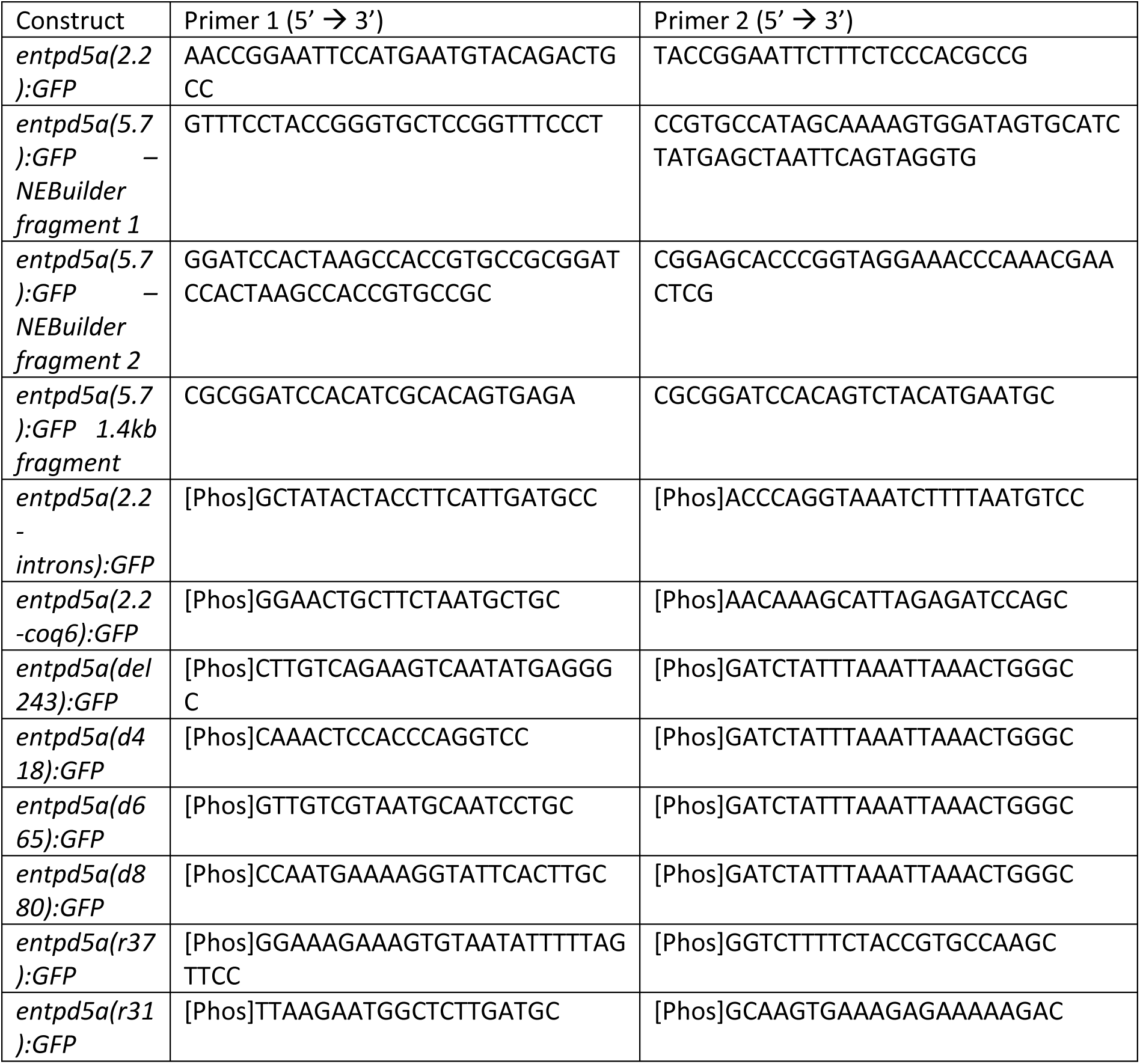
Primers used for cloning.

