## Supplementary figures and images for "Integration of ATAC and RNA-sequencing identifies chromatin and transcriptomic signatures in classical and non-classical zebrafish osteoblasts and indicates mechanisms of *entpd5a* regulation"

### SupplFig1

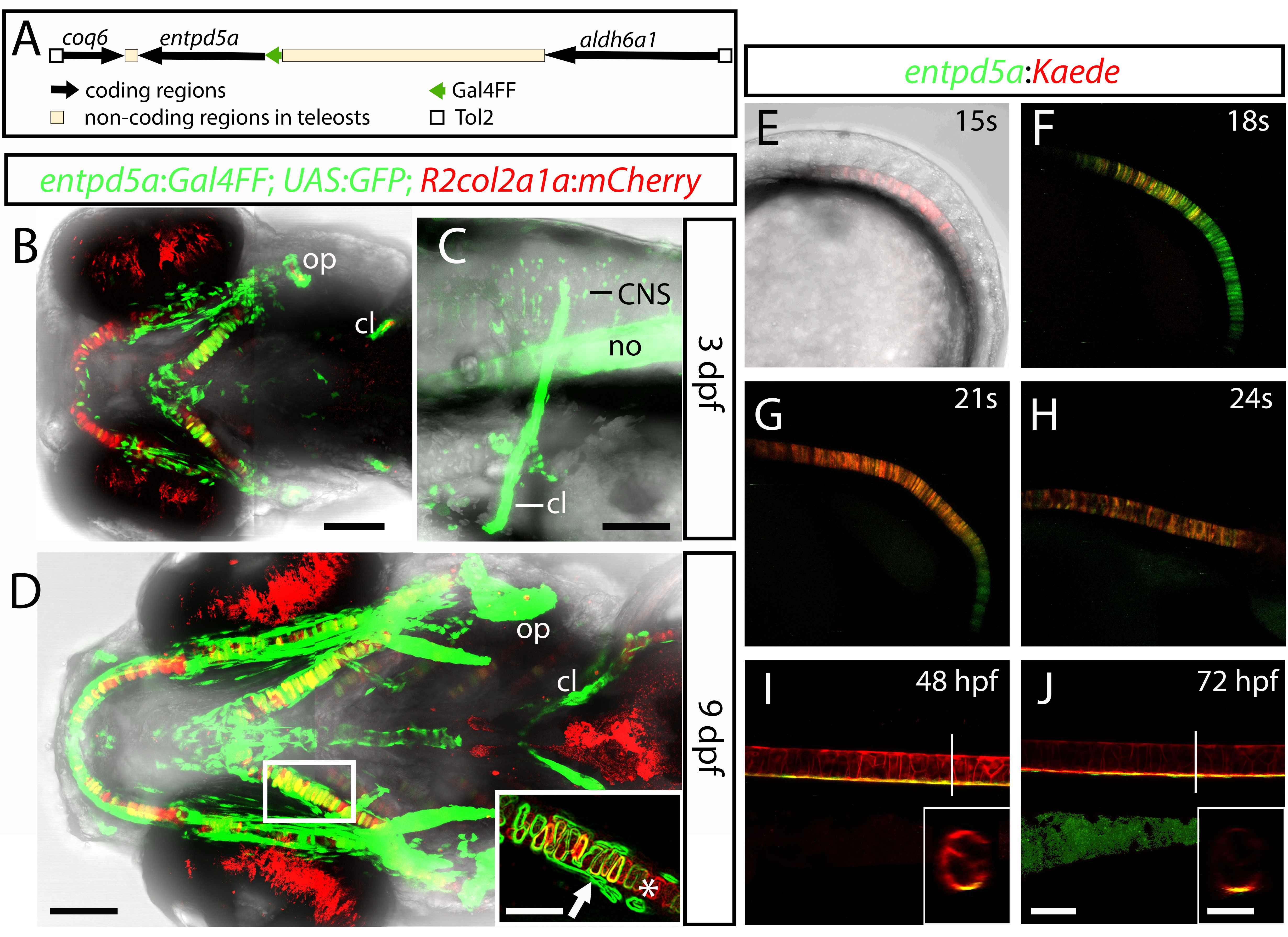

### SupplFig2

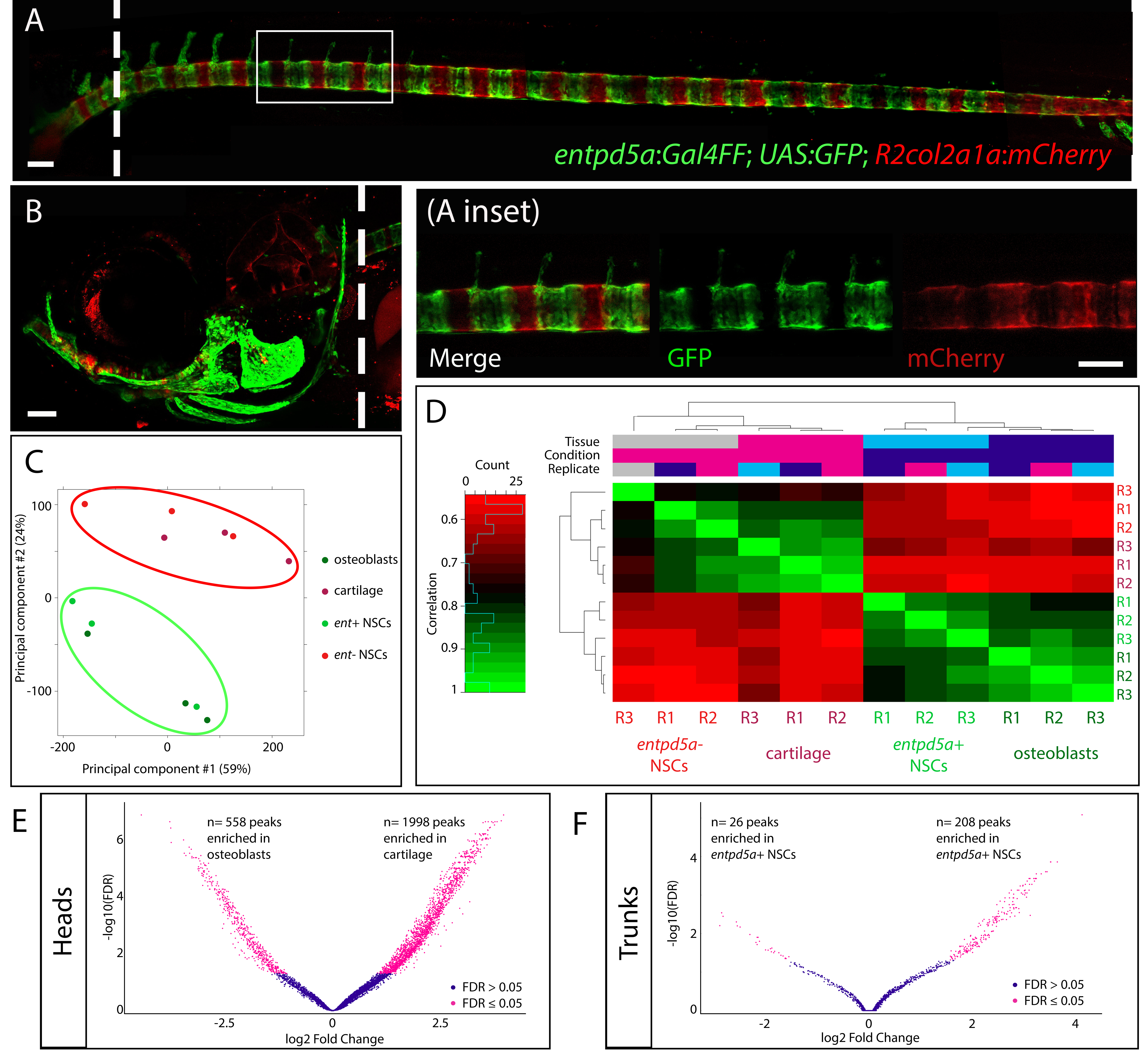

### SupplFig3

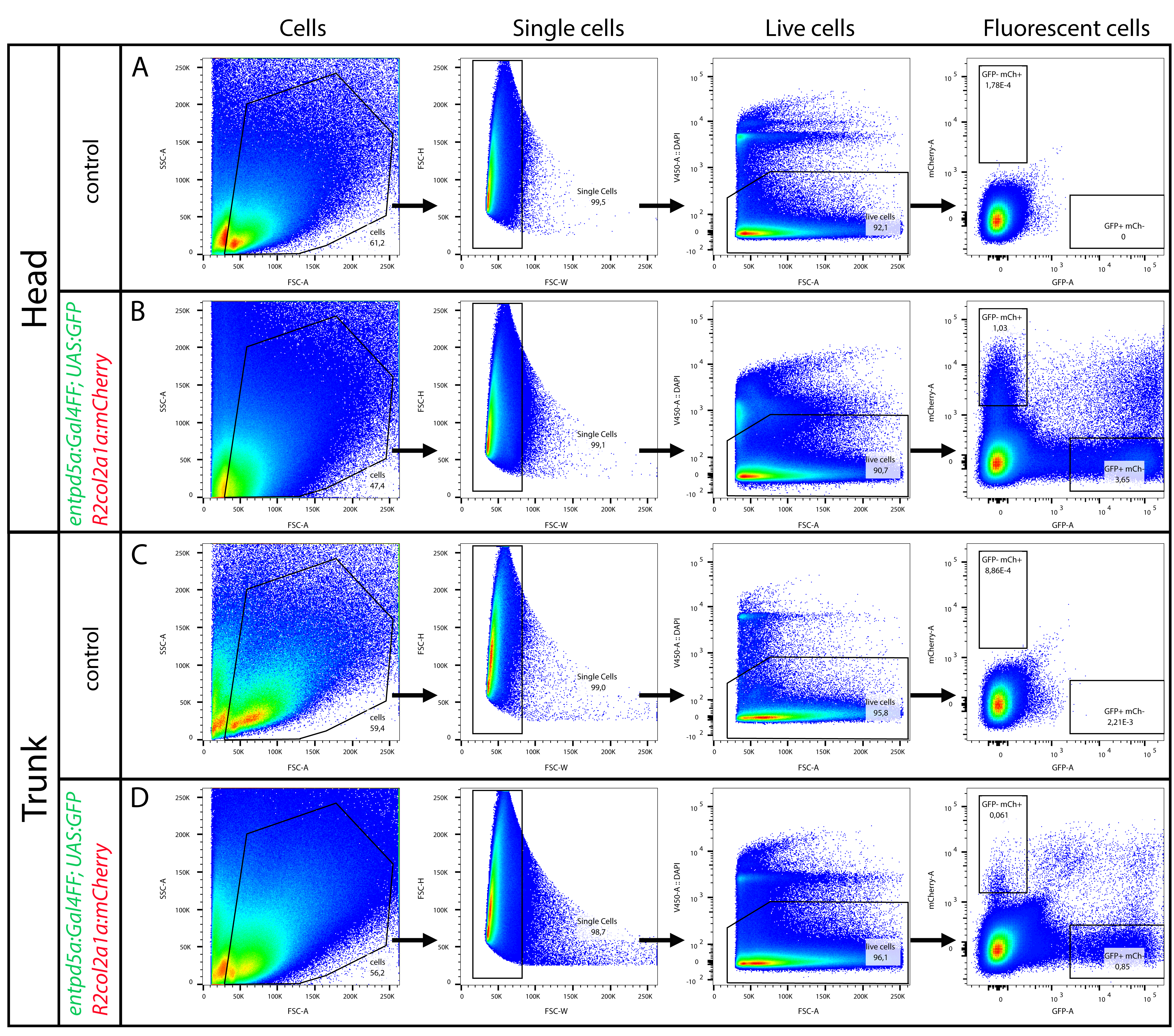

### SupplFig4

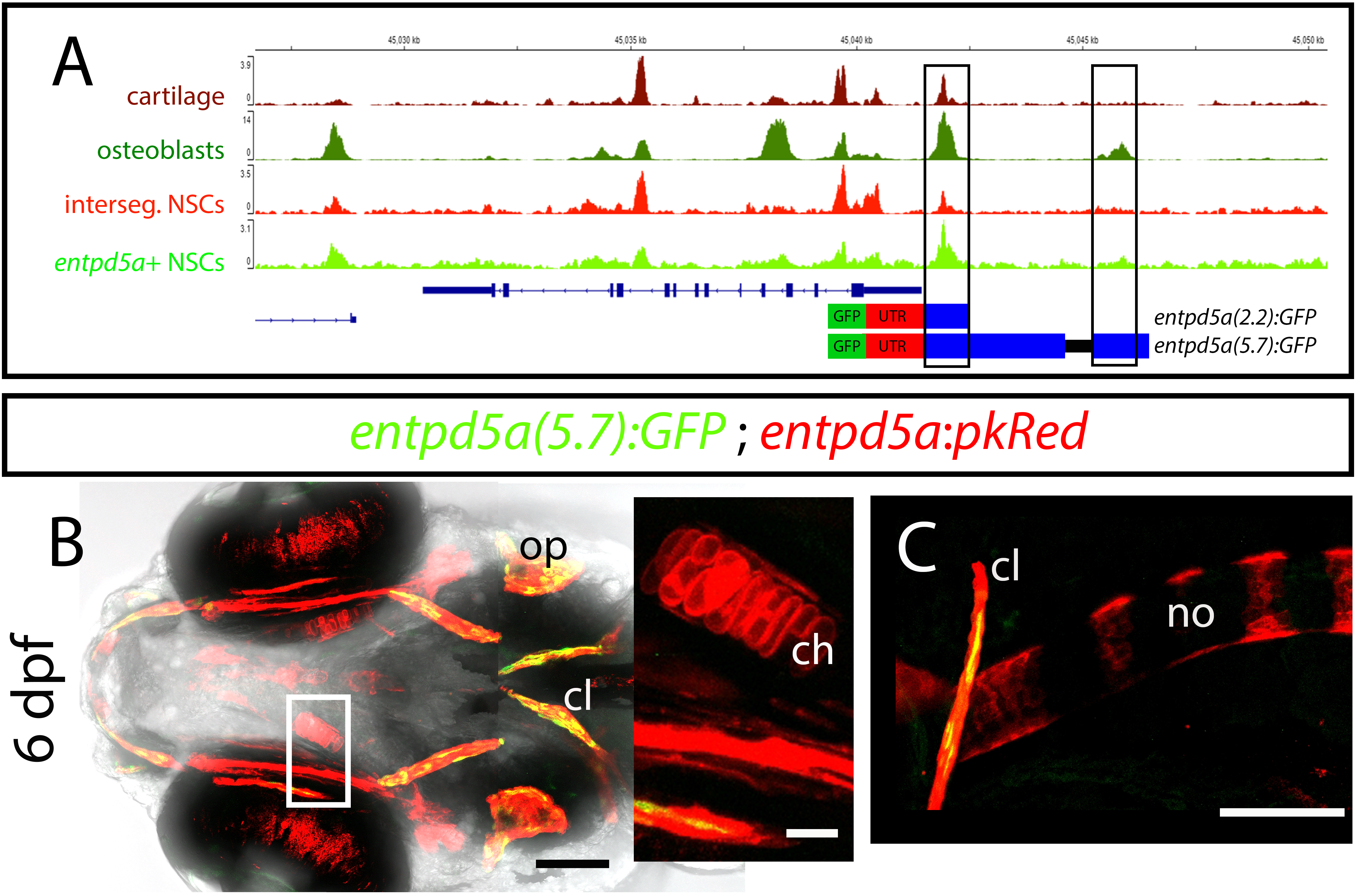

### SupplFig5

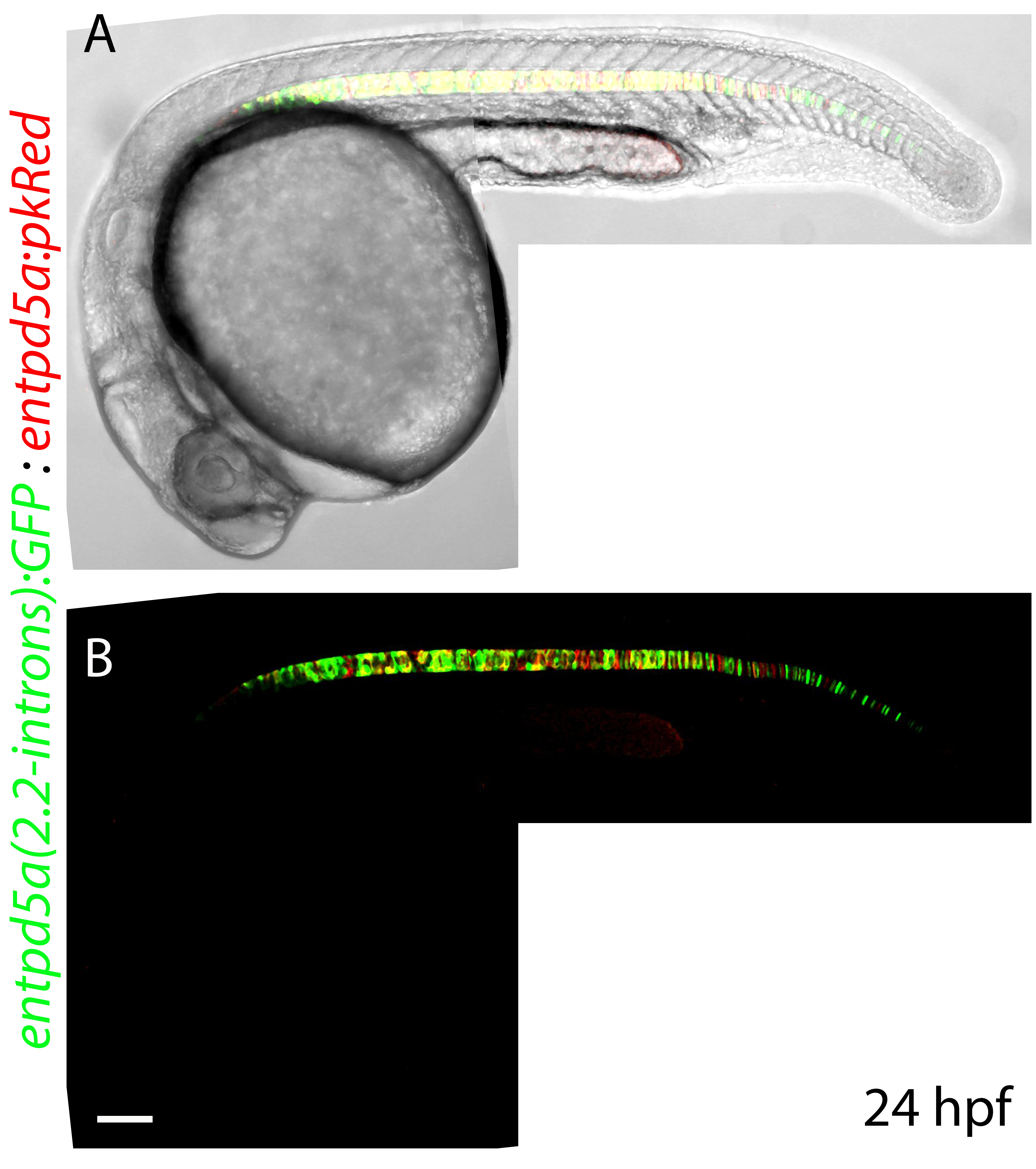

### SupplFig6

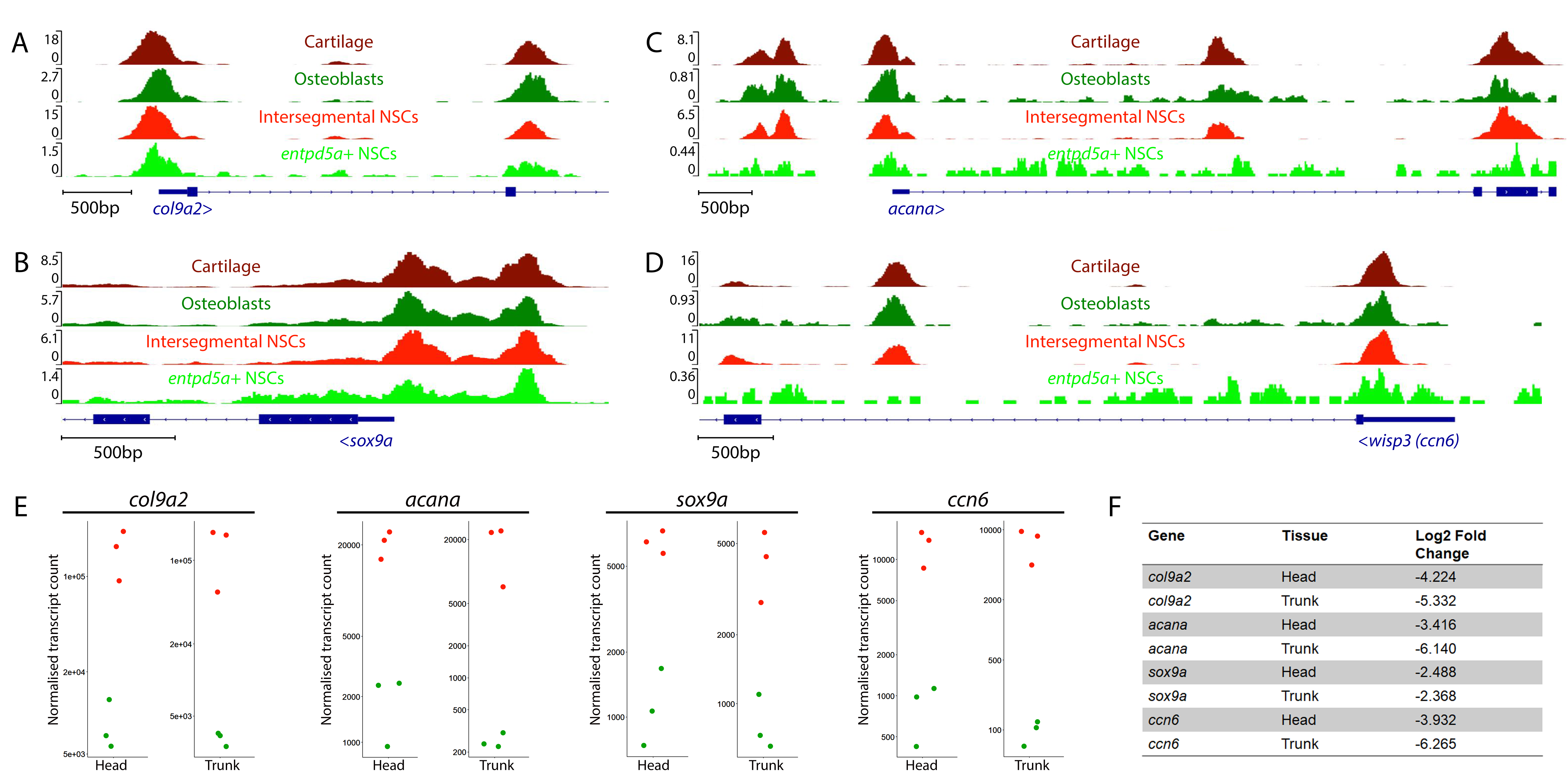
